# Preference for animate domain sounds in the fusiform gyrus of blind individuals is modulated by shape-action mapping

**DOI:** 10.1101/2020.06.20.162917

**Authors:** Łukasz Bola, Huichao Yang, Alfonso Caramazza, Yanchao Bi

**Author notes:** Corresponding authors: Łukasz Bola, Institute of Psychology, Polish Academy of Sciences, 1 Jaracza Street, 00-378 Warsaw, Poland, Yanchao Bi, National Key Laboratory of Cognitive Neuroscience and Learning & IDG/McGovern Institute for Brain Research, Beijing Normal University, Beijing 100875, China. These authors share first authorship (Łukasz Bola, Huichao Yang).

## Abstract

In high-level visual shape areas in the human brain, preference for inanimate objects is observed regardless of stimulation modality (visual/auditory/tactile) and subjects’ visual experience (sighted/blind individuals), whereas preference for animate entities seems robust only in the visual modality. Here, we test a hypothesis explaining this effect: visual shape representations can be reliably activated through different sensory modalities when they are systematically related to action system representations. We studied fMRI activations in congenitally blind and sighted subjects listening to animal, object, and human sounds. We found that, in blind individuals, the typical anatomical location of the fusiform face area responds to human facial expression sounds, with a clear mapping between the facial motor action and the resulting face shape, but not to speech or animal sounds. Using face areas’ activation in the blind subjects we could distinguish between specific facial expressions used in the study, but not between specific speech sounds. We conclude that auditory stimulation can reliably activate visual representations of those stimuli – inanimate or animate - for which shape and action computations are transparently related. Our study suggests that visual experience is not necessary for the development of functional preference for face-related information in the fusiform gyrus.

## Introduction

How does the brain transform modality specific signals into a representation that integrates stimulus properties from different senses? Here, we study an effect that might speak to this difficult question – that is, a stimulus domain by sensory modality interaction in high-level visual areas in the human occipitotemporal cortex (OTC).

The OTC hosts areas preferentially responding to stimuli from specific domains, such as manipulable inanimate objects that can be used as tools, stable inanimate objects relevant for navigation, and animate entities relevant for social interactions (Konkle and Caramazza, 2013; Magri, Konkle, and Caramazza, 2020). Interestingly, functional preference for inanimate objects in this region can be observed regardless of stimulation modality (visual/auditory/tactile) and subjects’ visual experience (sighted/blind individuals) (Mahon et al., 2009; Wolbers et al., 2011; He et al., 2013; Peelen et al., 2013; Dormal et al., 2018); in contrast, functional preference for animate entities seems considerably more tied to the visual modality, with a number of auditory and tactile studies reporting no preferential OTC responses for this domain (Goyal et al., 2006; He et al., 2013; Fairhall et al., 2014; Dormal et al., 2018). Recently, it has been proposed that this domain by sensory modality interaction in the OTC might reflect a difference in mapping between visual shape and action representations across inanimate and animate domains (Bi et al., 2016). This conjecture originates from a theoretical framework which assumes that the domain organization in the OTC is the result of evolutionary pressures to accommodate efficient mapping between locally-computed visual shape representations and appropriate downstream computations (Mahon & Caramazza, 2011). Within this framework, providing reliable information to the action system is a major function of the visual cortex, determining its organization. Critically, at the stage of the OTC, the relationships between visual and action representations are different across domains. In the case of inanimate objects, mapping between visual shape or texture and appropriate action representations is usually systematic and transparent (e.g., an elongation affords an action of gripping). This strong link between visual and action (i.e., non-visual) properties might make OTC representations of inanimate objects more accessible through different sensory systems. In the animate domain, the link between visual shape or texture and appropriate action representations is usually much less articulated (e.g., similarly looking animals or humans might or might not pose a threat). Lack of systematic relationship between shape and action representations may have resulted in OTC representations of animate entities being, on average, less accessible through other sensory systems.

Can the principle of mapping between visual shape and action representations be used to identify specific types of animate entities for which the OTC representation is actually reliably responsive to non-visual stimulation? In the present study, we address this question by investigating responses of the fusiform face area (FFA; Kanwisher et al., 1997; Kanwisher and Yovel, 2006) to face-related sounds. Several previous studies have already investigated whether the FFA shows functional preference for tactile, auditory, or imagined information about the human face, relative to other types of stimulation in these modalities. When sighted subjects are explicitly asked to imagine faces that they know or that were shown to them before the experiment, the preferential response of the FFA, relative to other imagery conditions, has usually been found (Ishai et al., 2000; O’Craven and Kanwisher, 2000; Goyal et al., 2006). However, surprisingly mixed results have been observed in sighted subjects not engaged in explicit imagery tasks and in congenitally and early blind subjects. Thus, no preferential FFA activation was reported when sighted and early blind subjects were listening to human voices pronouncing vowels (Dormal et al., 2018), when sighted and congenitally blind subjects were perceiving faces through vision-to-auditory sensory substitution devices (Plaza et al., 2015; no effects in the right hemisphere, a positive effect for blind subjects in the left counterpart of the FFA), or when such subjects were touching 3-D models of faces (Goyal et al., 2006; a positive effect was found for late blind subjects; Kitada et al., 2013). Furthermore, a null result for the FFA was reported when sighted subjects were making semantic decisions about famous people, based on a word cue (Fairhall et al., 2014). In apparent contrast to these reports, other studies showed preferential activation of the FFA when sighted and congenitally blind subjects were tactually identifying and comparing 3-D models of faces (Kitada et al., 2009; Murty et al., 2020). In the auditory modality, preferential FFA response was found in congenitally blind subjects listening to sounds of human facial actions (van den Hurk et al., 2017; repliacted in Murty et al., 2020) or to verbal phrases conveying different emotional states (Fairhall et al., 2017). While differences between the tactile studies might perhaps be explained by differences in statistical power (n = 3 per subject group in Goyal et al., 2006, where an effect in the FFA of congenitally blind and sighted individuals was not reported; a very subtle effect in the visual counterpart of the tactile experiment in Kitada et al., 2013), this methodological argument is not easily applicable to other studies discussed above.

Here, we propose a theoretical explanation of these apparently conflicting results. Specifically, in an extension of the shape-action mapping conjecture, we hypothesize that FFA responses to non-visual stimulation is modulated by the relationship between face shape and facial motor action representations. This view predicts that dynamic representations of face shapes, which are systematically and transparently related to specific types of facial motor actions, are more accessible through non-visual modality than other, more static face representations. While this modulatory effect might be more subtle in tactile or imagery studies, which encourage generation of shape descriptions (e.g., elongations or curvatures) that naturally map onto similar visual descriptions, independently of shape-action mapping, it should be clearly detectable in studies using sounds or words. In line with this hypothesis, previous auditory studies reported functional preference of the FFA for sounds of facial actions (van den Hurk et al., 2017) or phrases being indicative of the emotional state of a speaker (Fairhall et al., 2017) – that is, dynamic stimuli with a clear mapping between the facial motor action (e.g., an action of laughing) and the resulting face shape. In contrast, previous studies which did not observe functional preference of the FFA used stimuli for which such a relationship might be less clear, that is, speech sounds produced in a neutral tone (Dormal et al, 2018) or names of famous people (Fairhall et al., 2014).

To experimentally test our hypothesis, we designed a functional magnetic resonance imaging (fMRI) study in which congenitally blind and sighted subjects listened to object sounds and four categories of sounds produced by animate entities. The first two animate categories included human sounds associated with either emotional or non-emotional facial expressions (sounds of laughing and crying vs. sounds of yawning and sneezing). The other two animate categories included speech sounds produced in a neutral tone and animal sounds. As discussed above, the facial expressions are dynamic stimuli with a clear mapping between a facial motor action and a resulting face shape. This mapping can be learnt even in the absence of visual experience, through proprioception. We thus predict that, in both sighted and blind subjects, the FFA shows functional preference for both emotional and non-emotional facial expression sounds – a result that would allow us to distinguish between the FFA sensitivity to specific types of face-related information and sensitivity to emotional content, particularly in blind individuals (e.g., Fairhall et al., 2017). In contrast, we do not predict robust functional preference, neither in the FFA nor in the other ventral OTC (vOTC) animate areas, for speech and animal sounds.

## Methods

### Subjects

Twenty congenitally blind subjects (13 males, 7 females, mean age ± SD = 46.25 ± 13.34 y, average length of education ± SD = 9.4 ± 3.9 y) and 22 sighted subjects (15 males, 7 females, mean age ± SD = 45.91 ± 10.41 y, average length of education ± SD = 9.41 ± 2.09 y) participated in the study. The groups were matched for age, gender, handedness, and years of education (all p > 0.25). All subjects were native Mandarin speakers and had no history of neurological disorders. All sighted subjects had normal or corrected-to-normal vision. Detailed characteristics of each blind subject are provided in Table 1. All experimental protocols were approved by institutional review board of Department of Psychology Peking University, China (2015/05/04), as well as by the institutional review board of Harvard University (IRB15-2149), in accordance with the Declaration of Helsinki. Informed consent was obtained from each participant prior to study.

**Table 1.** Characteristics of blind subjects.

| Subject | Age | Gender | Years of education | Handedness | Cause of blindness | Light perception |
| --- | --- | --- | --- | --- | --- | --- |
| CB1 | 47 | F | 12 | R | Congenital glaucoma | None |
| CB2 | 30 | M | 0 | R | Congenital anophthalmos | None |
| CB3 | 40 | M | 12 | L | Congenital microphthalmia | None |
| CB4 | 54 | F | 12 | R | Congenital cataracts and eyeball dysplasia | Faint |
| CB5 | 52 | F | 9 | R | Congenital glaucoma | None |
| CB6 | 61 | M | 12 | R | Congenital eyeball dysplasia | None |
| CB7 | 67 | M | 9 | R | Congenital glaucoma and leukoma | None |
| CB8 | 52 | M | 12 | L | Congenital microphthalmia | None |
| CB9 | 44 | M | 12 | R | Congenital leukoma | Faint |
| CB10 | 52 | M | 12 | R | Congenital glaucoma | None |
| CB11 | 27 | M | 12 | R | Congenital glaucoma | None |
| CB12 | 52 | M | 12 | L | Congenital microphthalmia, cataracts and leukoma | None |
| CB13 | 34 | F | 12 | R | Congenital microphthalmia | None |
| CB14 | 28 | M | 0 | R | Congenital dysplasia, amotio retinae, aphakia and cataract | Faint |
| CB15 | 60 | F | 9 | R | Congenital microphthalmia and anophthalmos | None |
| CB16 | 59 | F | 9 | R | Congenital eye's tumor disease and dysplasia | None |
| CB17 | 54 | F | 9 | R | Congenital microphthalmia | None |
| CB18 | 59 | M | 9 | R | Congenital<br>anophthalmos and<br>cataract | None |
| CB19 | 23 | M | 11 | R | Unknown | None |
| CB20 | 30 | M | 3 | R | Congenital glaucoma<br>and cataract | None |

### fMRI experiment

In the fMRI experiment, subjects were presented with sounds belonging to five categories: emotional facial expressions (laughing and crying), non-emotional facial expressions (sneezing and yawning), speech sounds (two pinyin syllables – “bo” and “de”. The syllables conveyed no clear semantic information and were repeated three times in a neutral tone), object sounds (church bells, sleigh bells, sound of a car driving, sound of traffic) and animal sounds (a dog, a horse, a rooster and a cow). Each human sound was produced by two actors (man and woman). Furthermore, each sound subcategory included three unique sound exemplars (e.g., three different exemplars of a male actor laughing or three different exemplars of a horse neighing). All sounds were 2-s long and matched for sound intensity (root mean square value: −20 dB). However, since the sounds belonged to different object domains (e.g., animal versus human sounds), they were expected to inevitably differ in terms of many other auditory properties (see Fig. 1 for spectrograms and waveforms). Importantly, we expected to find a clear difference in the FFA activation only between facial expression sounds and other sound categories, even though differences in auditory properties were clearly present across all categories – thus, the differences in auditory properties alone cannot account for the pattern of results expected in this study. Sounds for the fMRI experiment were chosen from a larger dataset based on behavioral ratings of their emotional content (or lack thereof) and recognizability of depicted actions and sound categories, obtained from separate groups of sighted Chinese subjects (see Table S1 for the ratings for the sounds used in the fMRI study; demographic data of the raters: 11 males, 11 females, mean age ± SD = 24.05 ± 2.84 y, mostly students at Beijing Normal University). All human sounds were recorded specifically for this study. Object and animal sounds were downloaded from the internet.

**Figure 1.**
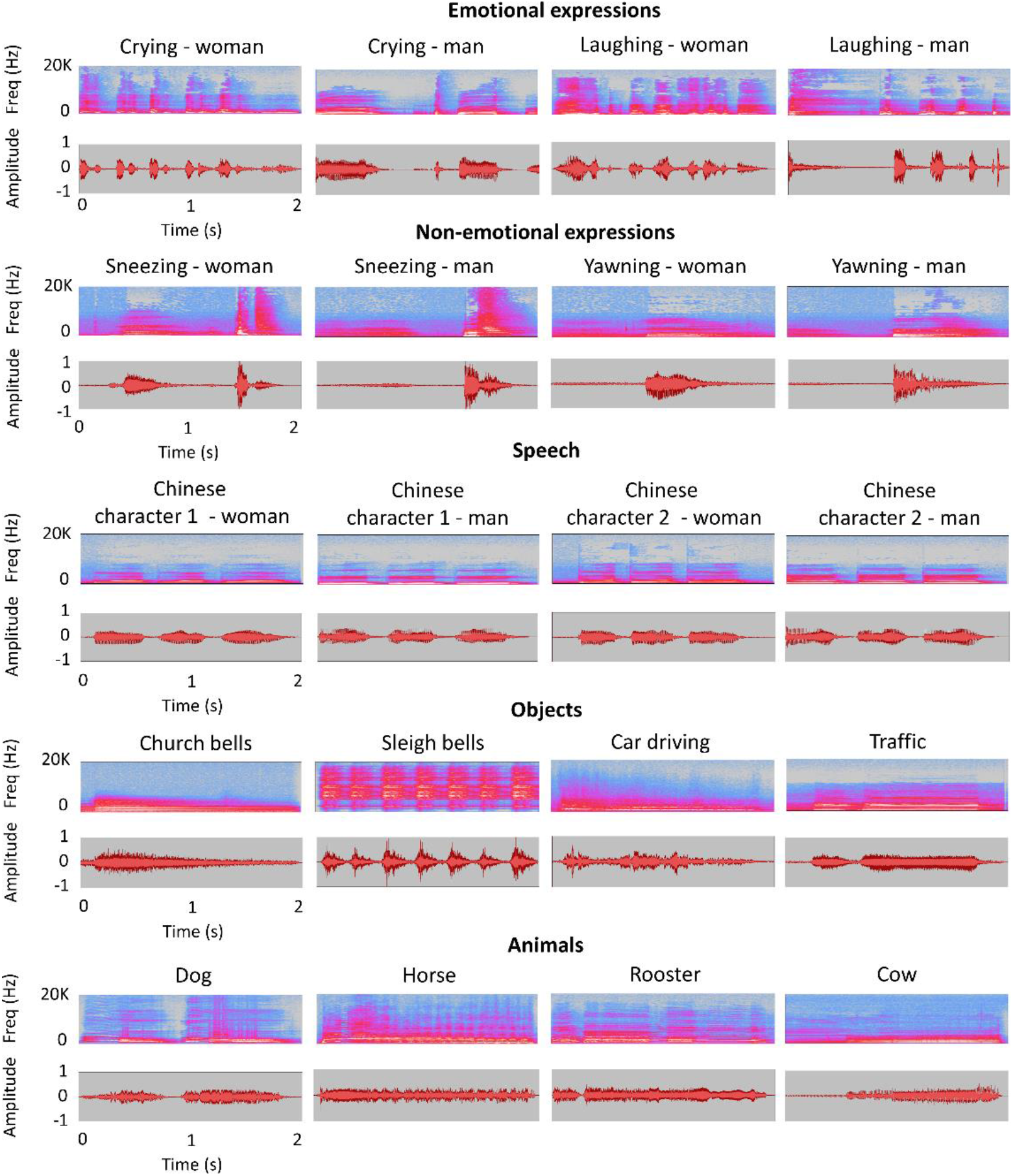
Spectrograms (upper panels) and waveforms (lower panels) of sounds used in the study. Each sound category included four subcategories, which, in turn, included three unique sound exemplars (e.g., three exemplars of a male actor laughing or three different exemplars of a horse neighing). The figure presents a representative exemplar of a sound from each subcategory. The spectrograms and the waveforms were produced using Audacity (https://www.audacityteam.org/).

The timing of the sound presentation in the fMRI scanner was kept the same across subject groups. Stimuli were presented in short blocks that included three sound exemplars belonging to the same condition and subcategory (e.g., three exemplars of church bell sounds). Presentation of each exemplar was followed by a 500-ms of silence period, resulting in a total duration of each block equal to 7.5 s. The order of specific sound exemplars’ presentation within each block was randomized in order to modulate purely auditory properties across blocks belonging to a given subcategory. The subjects’ task was to respond to each sound and answer whether it was produced by a human or non-human entity. For each sound exemplar, a response was recorded for 2.5 s following sound presentation’s onset.

Subjects participated in eight experimental runs. Each run included two blocks for every subcategory (e.g., male actor laughing), resulting in eight blocks for each category (e.g., emotional facial expressions) per run. Thus, 64 blocks for each category and 16 blocks for each subcategory were presented, in total. The blocks were interspersed with rest periods with no stimuli presented (from 0-to 18-s long) so that each run lasted 6.8 min. The order of each category presentation and the length of rest periods were determined using Optseq (https://surfer.nmr.mgh.harvard.edu/optseq/). A separate Optseq sequence was created for each run and the order of runs was randomized across subjects. Within each category of stimuli, the order of blocks belonging to specific subcategories was randomized with the rule that a given subcategory cannot be repeated two times in a row.

The stimulus presentation was controlled with a program written in Psychopy 3.0.12b (Peirce, 2007). The sounds were presented through MRI-compatible headphones. Subjects gave their answers by pressing buttons on a response pad with the index and the middle finger (e.g., the index finger – human sounds; the middle finger – non-human sounds), and a response mapping to a given finger was counterbalanced across subjects. Before the actual fMRI experiment, each subject completed a short training session outside the scanner, in order to ensure that the task was clear and that he/she can classify stimuli correctly.

### MRI data acquisition

Data were acquired on a Siemens Prisma 3-T scanner with a 20-channel phase-array head coil at the Imaging Center for MRI Research, Peking University. Functional data were acquired using simultaneous multi-slice sequence supplied by Siemens: slice planes scanned along the rectal gyrus, 62 slices, phase encoding direction from posterior to anterior; 2 mm thickness; 0.2 mm gap; multi-band factor = 2; TR = 2000 ms; TE = 30 ms; FA = 90°; matrix size = 112 × 112; FOV = 224 × 224 mm; voxel size = 2 × 2 × 2 mm. Right before the start of the collection of the first functional run, MRI field mapping was performed using the phase-magnitude sequence with following parameters: slice planes scanned along the rectal gyrus, 62 slices, phase encoding direction from posterior to anterior; 2 mm thickness; 0.2 mm gap; TR = 620 ms; TE^1^ = 4.92 ms; TE^2^ = 7.38 ms FA = 60°; matrix size = 112 × 112; FOV = 224 × 224 mm; voxel size = 2 × 2 × 2 mm. T1-weighted anatomical images were acquired using a 3D MPRAGE sequence: 192 sagittal slices; 1 mm thickness; TR = 2530 ms; TE = 2.98 ms; inversion time = 1100 ms; FA = 7°; FOV = 256 × 224 mm; voxel size = 0.5 × 0.5 × 1 mm; matrix size = 512 × 448.

### Behavioral data analysis

Accuracy obtained from the categorization task performed in the MRI scanner were entered into the group (blind subjects, sighted subjects) × sound category (emotional facial expressions, non-emotional facial expressions, speech, objects, animals) ANOVA. Analysis were performed in SPSS 25 (IBM). The reaction time was not analyzed since (1) the primary goal of the behavioral task was to verify whether subjects were attentive in the scanner, for which the accuracy is a better measure; (2) The time needed to categorize a given sound as belonging to a specific category is likely to depend not only on a subject effort or mental processes, but also on characteristics of the sound itself (e.g., its temporal dynamics). Thus, the reaction time is a rather uninformative measure in the context of our experiment.

### MRI data analysis

#### Data preprocessing

Before the actual preprocessing, the MRI data were converted to NIfTI format using dcm2niix (https://github.com/rordenlab/dcm2niix). Furthermore, anatomical images were deidentified using mri_deface script (https://surfer.nmr.mgh.harvard.edu/fswiki/mri_deface; Bischoff-Grethe et al., 2007) The preprocessing was performed using SPM 12 (https://www.fil.ion.ucl.ac.uk/spm/software/spm12/) and CONN 18b toolbox (www.nitrc.org/projects/conn; Whitfield-Gabrieli and Nieto-Castanon, 2012) running on MATLAB R2018b (MathWorks). Data from each subject were preprocessed using a standard CONN pipeline for volume-based analyses, which includes the following steps: (1) Realignment of all functional images to the first collected image; (2) Distortion correction and unwarping of functional images using a voxel-displacement map, created based on the field mapping sequence; (3) Slice-timing correction of functional images; (4) Detection of outliers in functional time series using the functions of Artifact Detection Tools toolbox (standard CONN “conservative” settings were used: all volumes showing, relative to the previous image, global signal change of z = 3 or subject’s head motion larger than 0.5 mm were marked as an outlier); (5) Coregistration of the anatomical image to the mean functional image; (6) Segmentation of the coregistered anatomical image and its normalization to Montreal Neurological Institute (MNI) space; (7) Normalization of all functional images to MNI space, using a deformation field obtained for the anatomical image; (8) Spatial smoothing with Gaussian kernel at two levels: 8 mm FWHM for the univariate analysis and 2 mm FWHM for the multivariate analyses (see Gardumi et al., 2016 for evidence that mild-to-moderate smoothing might improve the sensitivity of the decoding analysis).

Two first-level statistical models were created for each subject. For the univariate analysis, the data smoothed with 8-mm FWHM kernel were modeled at the level of sound categories (five experimental conditions: emotional expression sounds, non-emotional expression sounds, speech sounds, object sounds, animal sounds). For the multi-voxel pattern analysis (MVPA), the data smoothed with 2-mm FWHM kernel were modelled at the level of specific subcategories (twenty experimental conditions: five categories × four subcategories). In both models, the signal time course was modeled within a general linear model (Friston et al., 1995) derived by convolving a canonical hemodynamic response function with the time series of the experimental conditions. Additionally, six movement parameter regressors and a separate regressor for each volume identified as an outlier in the preprocessing were added to both models. An inclusive high-pass filter was used (400 s) to remove drifts from the signal while ensuring that effects specific to each subcategory are not filtered out from the data (note that only two blocks per subcategory were presented in each run and the order of presentation was randomized). Finally, individual beta estimates and contrast images were computed for each experimental condition (i.e., each category in the first model, each subcategory in the second model), relative to rest periods.

#### Univariate analysis

For the univariate whole-brain analysis, the contrast images for five sound categories vs. rest periods, for both groups, were entered into a random-effect 2 (Group) × 5 (Sound Category) ANOVA model, created using the SPM “flexible factorial” functionality. This model was used to perform all whole-brain univariate analyses reported, except for testing for the main effect of the group – since error terms calculated within the SPM flexible factorial design are generally not suitable for testing for main effects of the group, this analysis was performed separately, by entering individual average activations for all sound categories vs. rest periods into a simple two-group model (the SPM “two-sample t-test” functionality). F-tests were used to test for main effects of group and sound category and for the omnibus interaction between these factors. T-tests were used to perform a detailed pairwise comparisons. For each test at the group level, Threshold-Free Cluster Enhancement (TFCE; Smith and Nichols, 2009) values were calculated, with standard parameters (E = 0.5, H = 2; see Smith and Nichols, 2009), for every voxel within a broad grey matter mask, created based on canonical brain tissue probability maps implemented in SPM (all voxels in the brain image with a 20% or higher probability of belonging to the grey matter were included). The statistical significance of these values was then assessed using permutation testing (10000 iterations with labels of compared conditions randomized) and corrected for multiple comparisons using the false discovery rate (FDR; Benjamini and Hochberg, 1995). The calculation of these measures was performed using TFCE SPM toolbox (v. 1.77; http://www.neuro.uni-jena.de/tfce/). The results of group comparisons were visualized on a standard MNI152 template using MRIcroGL (https://www.nitrc.org/projects/mricrogl).

For the univariate region-of-interest (ROI) analysis, the FFA and the control ROIs - the primary auditory cortex (A1), the superior temporal sulcus (STS), and the part of the right posterior superior temporal sulcus responsive to facial expressions (right pSTS; Allison et al., 2000; Andrews et al., 2004; Pitcher et al., 2014) - were defined using a statistical association map produced by an automatic Neurosynth meta-analyses (v. 0.7; https://neurosynth.org/). The logic behind Neurosynth association maps is to highlight brain areas that are reported as activated in articles using a target term significantly more often than in articles that do not contain this term (Yarkoni et al., 2011). The FFA ROI was defined using a target term “face”. The meta-analysis included 896 studies and the association map was thresholded at Z-value = 15, resulting in a cluster in the right fusiform gyrus consisting of 227 voxels (functional data resolution; cluster center of mass MNI coordinates: 41 −49 −20). The A1, STS, and right pSTS control ROIs were defined using target terms “primary auditory” (114 studies; Z = 10; 672 voxels), “STS” (203 studies; Z = 7; 2234 voxels), and “facial expressions” (250 studies; Z = 4.8; 233 voxels), respectively. MarsBaR SPM toolbox (http://marsbar.sourceforge.net/) was used to extract an average contrast estimate for all voxels in a given ROI for each subject and experimental condition. Analogously to the univariate whole-brain analysis, these values were then entered into a random-effect 2 (Group) × 5 (Sound Category) ANOVA model, which was created in SPSS 25 (IBM). The results of pairwise comparisons were corrected for multiple comparisons using FDR. The FDR correction was calculated separately for three analysis steps – that is, for pairwise comparisons within each group (10 comparisons across sound categories per group) as well as for pairwise comparisons between groups (5 comparisons).

#### MVPA

The MVPA included both the classification (decoding) analysis and the representational similarity analysis (RSA). The FFA ROI was defined in a Neurosynth meta-analysis with a target word “face”, thresholded at Z-value = 10 (896 studies included). This resulted in a bilateral ROI consisting of 936 functional voxels (cluster center of mass MNI coordinates, right hemisphere: 42 −57 −18; anterior cluster in the left hemisphere: −41 −50 −19; posterior cluster in the left hemisphere: −39 −84 −13). The reason behind broader, compared to the univariate analysis, and bilateral ROI definition was that the MVPA relies on the disperse and subthreshold activation and deactivation patterns, which might be well represented also by cross-talk between hemispheres (for example, a certain subcategory might be represented by activation of the right FFA and deactivation of the left analog of this area). Performing MVPA in the FFA ROI used in the univariate analysis did not change the results in the blind group and yielded qualitatively similar, but statistically weaker results in the sighted group. The control ROIs in MVPA were the auditory cortex (A1 and STS univariate ROIs combined), the right pSTS (the Neurosynth meta-analysis with a target term “facial expressions”; 250 studies; Z = 2, 440 voxels), and the parahippocampal place area (PPA; Epstein and Kanwisher, 1998; the Neurosynth meta-analysis with a target term “place”; 189 studies; Z = 3, 790 voxels).

The MVPA was performed using CoSMoMVPA (v. 1.1.0; Oosterhof et al., 2016). The classification analysis was performed on single-subject beta estimates for each sound subcategory vs. rest periods using a support vector machine, as implemented in LIBSVM (v. 3.23; Chang and Lin, 2011). Cross-validation was performed using a leave-one-run-out procedure. A standard LIBSVM data normalization procedure (i.e., z-scoring beta estimates for each voxel in a test set and applying output values to the test set) was applied to the data before the classification. For the RSA, beta estimates were averaged across sound subcategories and runs to obtain one beta estimate for each of five sound categories, for a given subject. Correlation between the theoretical model of the FFA response, derived from our hypothesis, and actual responses in this area was performed using Spearman’s rank correlation coefficient.

Obtained classification accuracies and RSA correlation values were tested against chance level in a permutation analysis. Specifically, each analysis was rerun 1000 times for each subject with sound category labels randomly reassigned in each iteration. Null distributions created in this way were averaged across subjects, within each group, and compared with the mean classification accuracy obtained in the actual analysis. The FDR correction was applied to the results when appropriate (see figure legends for details).

## Results

### Behavioral results

To keep the subjects attentive in the MRI scanner, we asked them to categorize presented stimuli into human and non-human sounds. Both groups of subjects were highly accurate in performing this task (mean accuracy ± SD, blind subjects: 95% ± 6%; sighted subjects: 92% ± 10%). No difference between groups or sound categories were detected; the interaction between these two factors was also not significant (Group × Sound Category ANOVA, all p > 0.18).

### fMRI results: univariate analysis

We started the analysis of the fMRI data from performing omnibus F-tests, testing for main effects of sound category, group, and for interaction between these two factors. The main effect of sound category was observed in a wide network of regions, including frontoparietal regions, the temporal lobe, the OTC, and the early visual cortex (Table S2). When we increased the statistical threshold, in order to achieve a higher degree of spatial specificity, we detected the main effect of condition primarily in the auditory cortex, right frontal areas and the left cerebellum (Fig. S1). Testing for the main effect of group did not yield significant results. A group by condition interaction was detected primarily in visual areas, including the OTC, as well as in the left and the right pSTS (Fig. S1; Table S3). To disentangle effects induced in the OTC by specific sound categories, in each group, we proceeded with detailed pairwise comparisons.

#### Congenitally blind group

##### Animate sound categories vs. object sounds

We first contrasted activation for sounds of every animate category with activation for object sounds in the blind group. As expected, sounds of both emotional and non-emotional expressions induced stronger response, relative to object sounds, in classic face processing areas - the right pSTS, the right lateral fusiform gyrus (the typical anatomical locus of the FFA) and the right inferior occipital gyrus (the typical anatomical locus of the occipital face area – OFA; Gauthier et al., 2000) (Fig. 2A-B; see also Table S4-5). Furthermore, sounds of emotional expressions only induced stronger activation in the amygdala (Fig. 2A; peak MNI coordinates, left hemisphere = −26 −8 −20; right hemisphere = 22 −2 −24). Outside the canonical face perception network, stronger responses to both types of facial expression sounds were observed in the primary visual cortex, dorsolateral visual areas, the auditory cortex and frontal regions (primarily the precentral gyrus, the inferior frontal gyrus and the middle frontal gyrus) (Fig. 2A-B; Table S4-5). Contrasting the other animate categories – that is, speech sounds and animal sounds - with object sounds yielded significant effects in the auditory cortex, the precuneus, frontal regions and, in the case of animal sounds, in some parts of the primary visual cortex (Fig. S2; Table S6-7). However, in line with our hypothesis, no reliable activation in the vOTC animate areas was observed for these animate sound categories (Fig. S2; Table S6-7).

**Figure 2.**
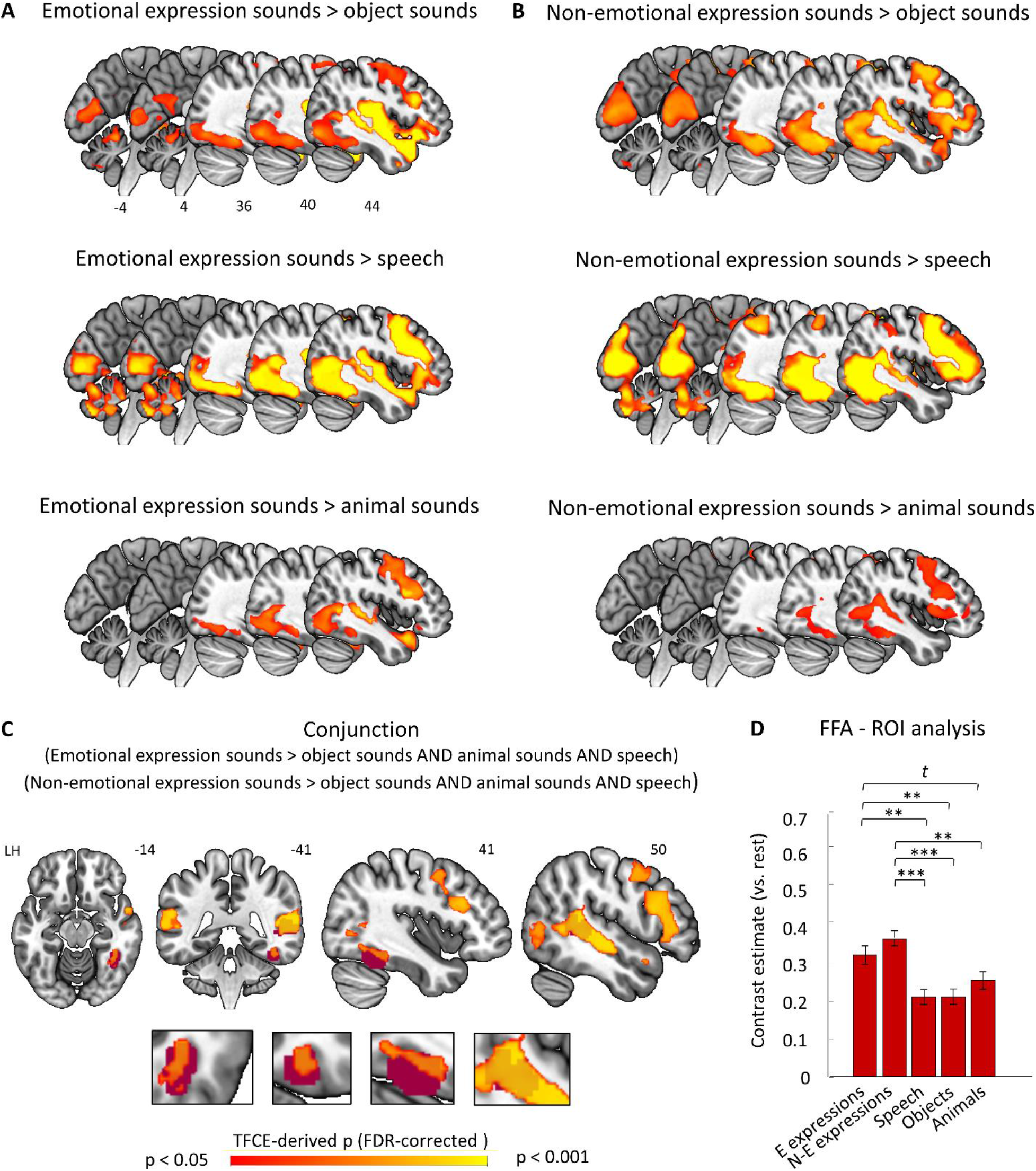
Functional selectivity for facial expression sounds in the fusiform gyrus of congenitally blind subjects. (A-B) Activation for the facial expression sounds relative to the activation for object sounds, speech sounds, and animal sounds. Results are presented separately for the expression sounds conveying emotional information (laughing and crying) and for the expression sounds that do not convey such information (yawning and sneezing). (C) Conjunction of the six contrasts presented in panels A and B. The burgundy ROIs represent the typical location of the FFA, as indicated by the Neurosynth meta-analysis with a target term “face” (896 studies included; map thresholded at Z = 15), and the typical location of the right pSTS responsive to facial expressions, as indicated by an analogous metanalysis with a target term “facial expressions” (250 studies included; map thresholded at Z = 4.8). Close-ups of the two ROIs, in the planes corresponding to the whole-brain illustrations, are provided at the bottom for clarity. (D) Mean contrast estimates for each sound category in the FFA ROI (E Expressions – emotional expressions; N-E expressions – non-emotional expressions). Statistical thresholds: (A-C) p < 0.05 derived through permutation of TFCE values and corrected for multiple comparisons using FDR. In the conjunction analysis, color hues represent mean FDR-corrected p-values of the six contrasts included. These values were calculated only for voxels that passed the conjunction test (i.e., were significant in each contrast included). (D) ** p < 0.01, *** p < 0.001, *t* = 0.09, FDR-corrected for multiple comparisons. Error bars represent standard error of the mean, adjusted to adequately represent within-subject variability across conditions using a method proposed by Cousineau (2005). Numbers in panels A and C denote MNI coordinates of presented axial, coronal or sagittal planes.

##### Facial expression sounds vs. other animate sound categories

We then directly compared activation enhanced by facial expression sounds with activation for other animate sound categories in the blind group. These comparisons confirmed that, relative to speech sounds and animal sounds, both types of the facial expression sounds induced stronger activation in the vOTC – particularly, in the typical anatomical location of the FFA and the OFA (Fig. 2A-B; Table S8-11). Stronger activation for facial expression sounds was also observed in dorsolateral visual areas, the right pSTS, the auditory cortex and frontal regions (Fig. 2A-B; Table S8-11). In the comparison with speech sounds, but not with animal sounds, stronger activation for both types of facial expression sounds was observed in the primary visual cortex (Fig. 2A-B; Table S8-11).

##### Facial expression sounds – functional selectivity analysis

To statistically test for areas showing functional selectivity for the facial expression sounds, relative to all other sounds used in the experiment, in the blind subjects, we performed a conjunction (i.e., logical AND) analysis of all contrasts between the sounds of facial expressions and the other sound categories (six contrasts included, see Fig. 2C). This stringent test of our hypothesis yielded a significant effect in the right lateral fusiform gyrus, in the typical location of the FFA (cluster center of mass MNI coordinates: 41 −48 −15; Fig. 2C). Furthermore, significant conjunction effects were also observed in the dorsolateral parts of the right visual cortex (the V5/MT area), the right STS, including the right pSTS, the left pSTS and right frontal regions (Fig. 2C).

#### Sighted group

##### Animate sound categories vs. object sounds

In the sighted group, both types of the facial expression sounds induced stronger response, relative to object sounds, in the auditory cortex and the pSTS, similarly to what was observed in the blind subjects (Fig. 3A-B; Table S12-13). Preferential activation for the facial expression sounds was also detected in frontal regions, although in the sighted subjects this effect was constrained primarily to the inferior frontal gyrus (Fig. 3A-B; Table S12-13). Intriguingly, and in contrast to the results for the blind group, no preferential response in the fusiform gyrus was observed in the sighted subjects, neither for the emotional nor for the non-emotional facial expression sounds (Fig. 3A-B; Table S12-13). The emotional facial expression sounds induced some preferential activations, relative to object sounds, in other visual regions – particularly in the dorsolateral visual areas, the posterior part of the inferior occipital gyrus and the cuneus (Fig. 3A; Table S12). However, similar effects were also observed in comparisons between speech sounds and object sounds, and, in the case of cuneus, between animal sounds and object sounds (see Fig. S3; Table S14-15).

**Figure 3.**
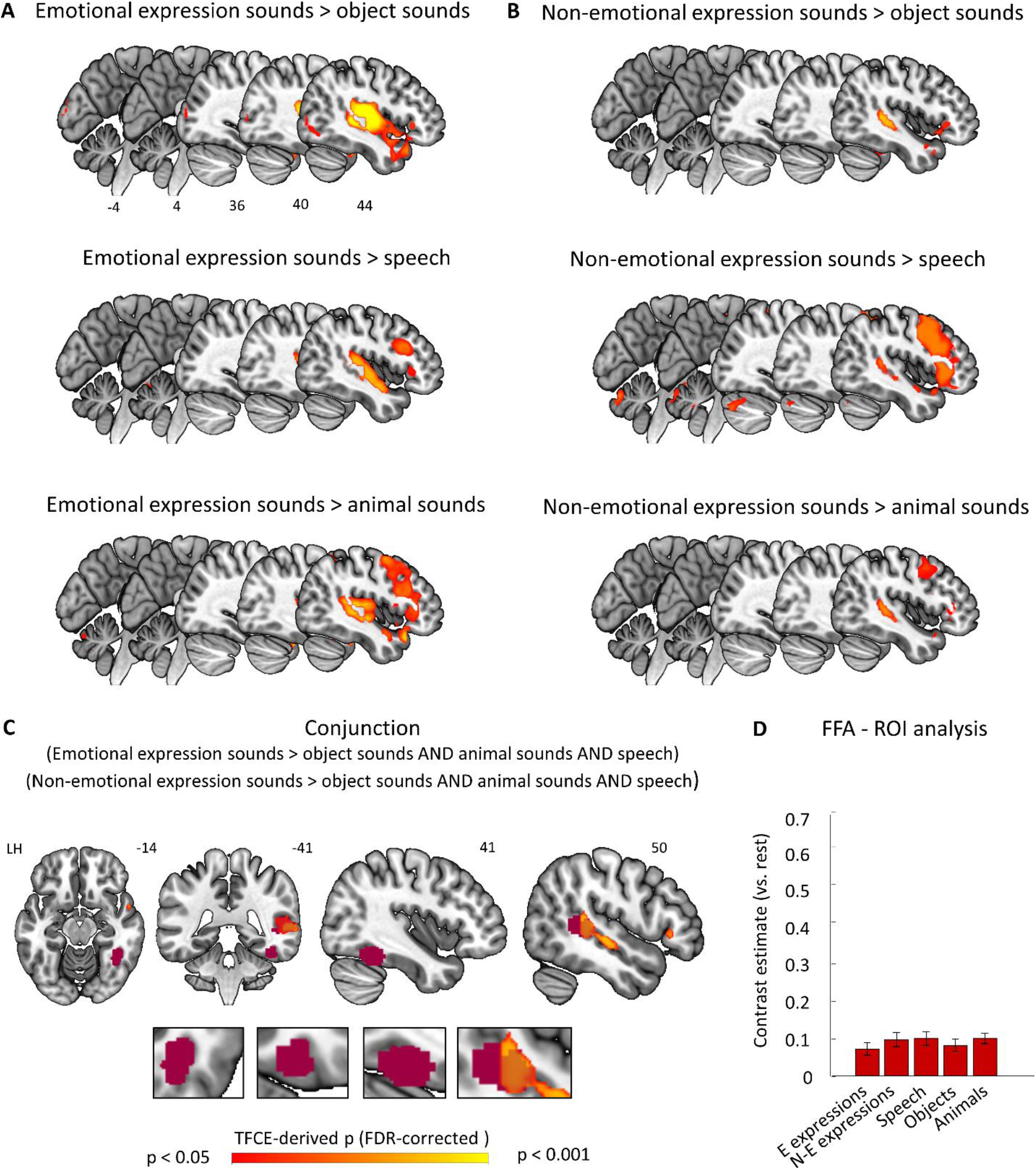
Lack of preferential univariate activation for facial expression sounds in the fusiform gyrus of sighted subjects. All analysis parameters and ROI definition procedures were identical to those applied to the data from the blind group (see Figure 2 for details). (A-B) Activation for the facial expression sounds relative to the activation for object sounds, speech, and animal sounds. Results are presented separately for the expression sounds conveying emotional information (laughing and crying) and for the expression sounds that do not convey such information (yawning and sneezing). (C) Conjunction of the six contrasts presented in panels A and B. The burgundy ROIs represent the typical anatomical location of the FFA and the part of the right pSTS that is responsive to facial expressions, as determined by Neurosynth meta-analyses for the words “face” and “facial expressions”, respectively (see Figure 2 for details). Close-ups of the two ROIs, in the planes corresponding to the whole-brain illustrations, are provided at the bottom for clarity. (D) Mean contrast estimates for each sound category in the FFA ROI (E Expressions – emotional expressions; N-E expressions – non-emotional expressions).

##### Facial expression sounds vs. other animate sound categories

Direct comparisons between the facial expression sounds and the other animate sound categories in the sighted subjects showed significant effects in the auditory cortex, the right pSTS and frontal regions; however, no significant differences were detected in the visual cortex (Fig. 3A-B; Table S16-19).

##### Facial expression sounds – functional selectivity analysis

The conjunction analysis of all six contrasts between the facial expression sounds and the other sound categories in the sighted subjects yielded significant effects in the left and the right auditory cortex, the left and the right STS, including the right pSTS, and the right inferior frontal gyrus; however, it did not show any significant effects in the visual cortex (Fig. 3C).

#### Between-group comparisons and ROI analyses

##### Between-group differences in sensitivity to facial expression sounds

To directly test for differences in functional preference for the facial expression sounds across groups, we performed an additional condition (average from all facial expression sounds > average from all other sounds) by group interaction analysis. In line with the results observed within each group, this analysis indicated a stronger preference for facial expression sounds in the right lateral fusiform gyrus and in the right inferior occipital gyrus (typical anatomical locations of the FFA and the OFA, respectively) in the blind subjects (Fig. 4; Table S20). Furthermore, the same effect was observed in the early visual cortex and in the dorsolateral visual areas, bilaterally, as well as in the most posterior parts of the left and the right pSTS (Fig. 4; Table S20). An inverse contrast, testing for brain areas showing stronger functional preference for facial expression sounds in the sighted group, did not yield any significant results.

**Figure 4.**
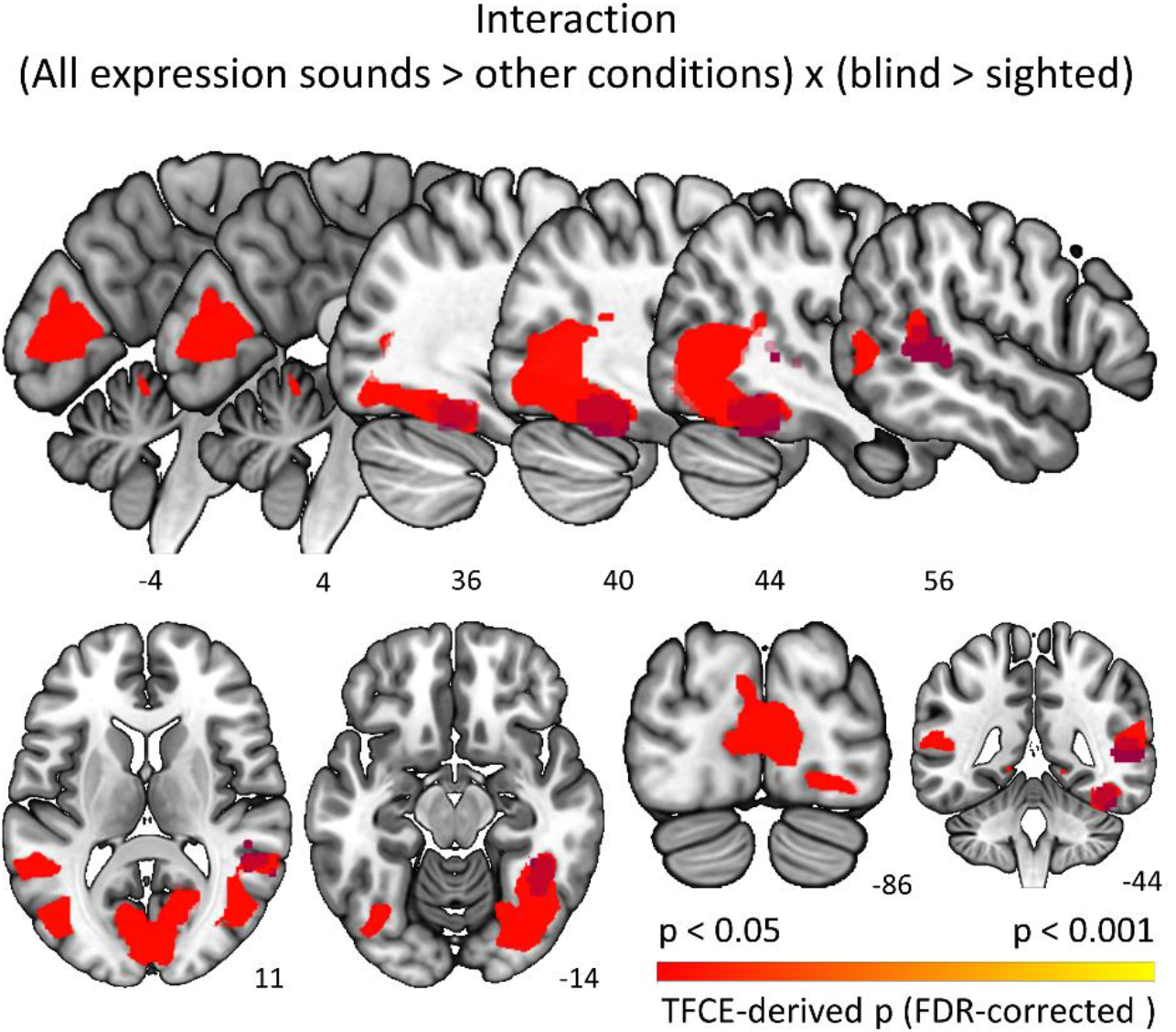
Direct test of between-group differences in sensitivity to facial expression sounds. Brain regions preferentially activated by facial expression sounds (averaged activation for emotional and non-emotional expressions), relative to other experimental conditions (averaged activation for speech, object sounds, and animals sounds), in the blind subjects but not (or not to the same extent) in the sighted subjects. All analysis parameters and ROI definition procedures were identical to those that were applied to within-group comparisons (see Figure 2 for details). The burgundy ROIs represent the typical anatomical location of the FFA and the part of the right pSTS that is responsive to facial expressions, as determined by Neurosynth meta-analyses for the words “face” and “facial expressions”, respectively (see Figure 2 for details). Numbers denote MNI coordinates of presented axial, coronal or sagittal planes.

##### ROI analysis in the FFA

To further confirm spatial correspondence between the effect observed for facial expression sounds in the blind subjects and the typical anatomical locus of the FFA, we performed an ROI analysis, in which the FFA ROI was defined based on the meta-analysis of face perception studies (see Methods). In a 2 (Group) × 5 (Sound Category) ANOVA, we detected significant main effects of sound category (F(4, 160) = 5.02, p = 0.001, partial eta squared = 0.11) and group (F(1, 40) = 6.87, p = 0.012, partial eta squared = 0.15) as well as significant interaction between these factors (F(4, 160) = 5.65, p < 0.001, partial eta squared = 0.12). In line with whole-brain analysis results, pairwise comparisons showed stronger activation for both types of facial expressions sounds than for any other sound category included in the study in the blind group (Fig. 2D). In contrast, no significant differences between conditions were detected in the sighted group (Fig. 3D). Furthermore, direct comparisons between groups showed stronger activation in the blind group for emotional facial expressions (mean difference = 0.25, SEM = 0.08, p_FDR_ = 0.01) and non-emotional facial expressions (mean difference = 0.27, SEM = 0.08, p_FDR_ = 0.005). Trends towards a similar difference for other sound categories - animal sounds (mean difference = 0.16, SEM = 0.08, p_FDR_ = 0.075), speech sounds (mean difference = 0.11, SEM = 0.07, p_FDR_ = 0.1) and object sounds (mean difference = 0.13, SEM = 0.07, p_FDR_ = 0.1) – were also detected.

##### ROI analysis in the right pSTS

The whole-brain analysis suggests that the functional preference for the facial expression sounds used in our study can be observed, in both groups, in the part of the right pSTS that is known to be sensitive to information about dynamic features of face shape (Allison et al., 2000; Andrews et al., 2004; Pitcher et al., 2014). To confirm this important control result, we performed an ROI analysis, in which the right pSTS was defined based on the meta-analysis of studies investigating brain responses to facial expressions (see Methods). The 2 (Group) × 5 (Sound Category) ANOVA showed a significant main effect of sound category (F(4, 160) = 35.91, p < 0.001, partial eta squared = 0.47), whereas the main effect of group and the interaction between the two factors were not significant (all p > 0.15). As expected, in both groups, post hoc tests showed stronger activation for both types of facial expression sounds relative to all other sound categories used in the experiment (Fig. S4; all p < 0.01). Thus, while the whole-brain interaction analysis hinted at stronger functional preference for facial expression sounds in the most posterior part of the right pSTS in the blind group (Fig. 4), an activation pattern averaged from this whole area is comparable across groups.

##### ROI analysis in auditory cortices

The sounds from different domains and semantic categories inevitably differ on many auditory properties (see Methods). Could differences in acoustic features or acoustic complexity drive the fusiform gyrus response pattern in the blind subjects? We find this possibility highly unlikely as the difference in activation of this region was found only in comparison between facial expression sounds and other sound categories. We did not observe activation differences in comparisons between other sounds categories (for example, between object sounds and animal sounds), despite the fact that these categories were also markedly different in terms of their auditory properties. To illustrate this point further, we performed the ROI analyses in the A1 and in the STS, which hosts higher-level auditory areas (Fig. S5 and S6). In both these ROIs, the activation patterns were similar in the blind and the sighted group and markedly different from the pattern observed in the fusiform gyrus in the blind group. This finding confirms that differences in auditory properties cannot explain the results reported here.

### fMRI results: MVPA

#### Multi-voxel pattern classification of sound categories

The univariate analysis of the fMRI data revealed strong functional selectivity to facial expression sounds in the typical anatomical location of the FFA in the blind subjects. In contrast, this area did not show any univariate differentiation of presented sounds in the sighted group. Nonetheless, it is still possible that the FFA in the sighted subjects represents the difference between the facial expression sounds and the other sound categories by the means of disperse, subthreshold activation patterns. To investigate this possibility, we performed a multi-voxel pattern classification in this area (see Methods).

Analogously to the univariate analysis, we first asked which categories of animate sounds can be distinguished from object sounds, based on the lateral fusiform gyrus activation patterns in each group. We found that a classifier, trained on the activation of this area, was able to successfully differentiate between both types of expression sounds and object sounds, in both blind and sighted subjects (Fig. 5A-B). Successful classification of sounds was also achieved for the speech - object sound pair (blind subjects: mean classification accuracy = 62 %, SEM = 2.4 %, p_FDR_ < 0.001; sighted subjects: mean classification accuracy = 55 %, SEM = 1.3 %, p_FDR_ = 0.001) but not for the animal - object sound pair (blind subjects: mean classification accuracy = 52 %, SEM = 1.6 %, p_FDR_ = 0.093; sighted subjects: mean classification accuracy = 52 %, SEM = 1.4 %, p_FDR_ = 0.093).

**Figure 5.**
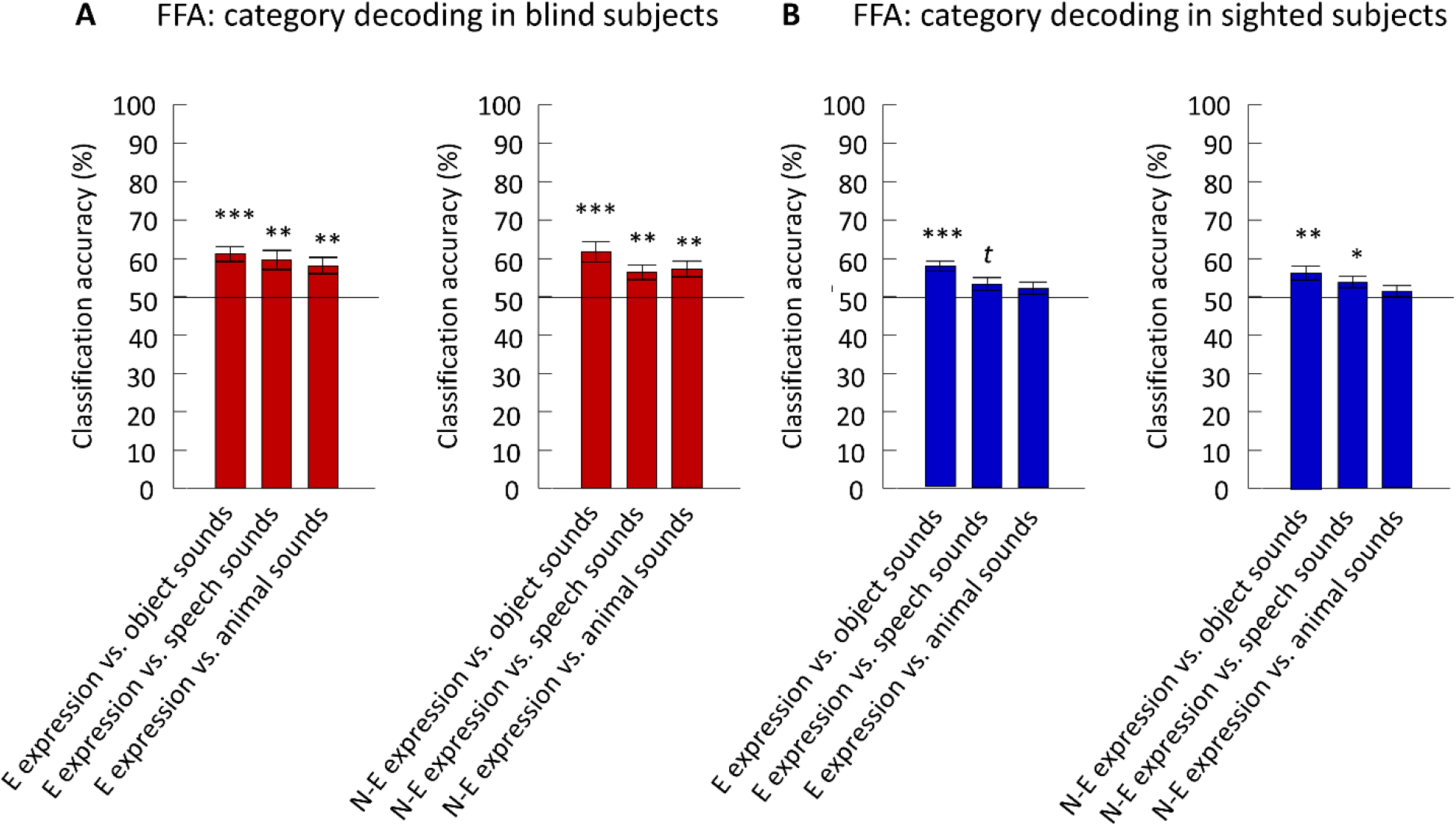
Facial expression sounds induce distinctive multi-voxel activation patterns in the fusiform gyrus, in both blind and sighted subjects. (A-B) Mean accuracy of a support vector machine classifier distinguishing between facial expression sounds and every other sound category included in the experiment, based on the activation patterns in the FFA. Results are presented separately for the blind (A) and the sighted (B) subjects and for the emotional expression sounds (E expression sounds; left panels) and the non-emotional expression sounds (N-E expression sounds, right panels). The FFA was defined based on the Neurosynth meta-analysis for the term “face” (896 studies included; the resulting map thresholded at Z = 10; for the purpose of the classification analysis, voxels in both the right and the left hemisphere were included, see Methods). Testing against chance was performed in a permutation analysis. Statistical thresholds: * p < 0.05, ** p < 0.01, *** p < 0.001, *t* p = 0.057, corrected for multiple comparisons across all tests performed using FDR. Error bars represent standard error of the mean. Chance classification level is marked with a black line.

We then tested whether the activation of the lateral fusiform gyrus can be used to distinguish between facial expression sounds and other animate sound categories. Indeed, we observed a successful classification of sounds into facial expression sounds and speech sounds, in both groups and in the case of both types of facial expressions (Fig. 5A-B). Additionally, the classifier was able to distinguish between the facial expression sounds and animal sounds, in the blind, but not in the sighted subjects (Fig. 5A-B).

We repeated the same analyses in three control ROIs - the broadly-defined auditory cortex (the primary auditory cortex and the STS ROIs combined), the PPA (Epstein and Kanwisher, 1998), and the right pSTS. We documented highly successful classification of all category pairs, in both groups, in the auditory cortex (a positive control site; Fig. S7) and in the right pSTS (Fig. S8). In contrast, classification of animate category pairs was generally not successful in the PPA (a negative control site; Fig. S9). As can be expected, this area seemed to distinguish mostly between the object sounds and the animate sound categories – especially in the blind group (Fig. S9).

#### Multi-voxel pattern classification of specific facial expression sounds

Does the typical anatomical location of the FFA code the differences between specific facial expressions presented through the auditory modality? To address this question, we trained a classifier to distinguish, based on the activation of this area, between four facial expressions presented in our study (crying, laughing, sneezing and yawning), irrespectively of the differences in voice characteristics and in the gender of the two actors (man and woman; see Methods). Indeed, the sounds were successfully classified into specific facial expressions in the blind group (Fig. 6A). The confusion matrix illustrating classifier performance (i.e., its choices for specific stimulus classes) showed that successful classification was not driven by only one specific sound category or by purely categorical difference between emotional or non-emotional expression sounds (Fig. 6B). Contrary to the result for facial expression sounds, no significant effects were observed in the blind group for the classification of speech sounds, irrespectively of the gender of the two speakers, or for the classification of the gender of the two actors/speakers, irrespectively of the specific facial expression or speech sound (Fig. 6A). In the sighted group, none of the above-described analyses yielded significant results (Fig. 6A). The control analysis in the auditory cortex yielded a successful classification for all conditions and both groups (Fig S10). In contrast, none of the analyses performed were significant in the PPA (Fig S11). Finally, the activation of the right pSTS was diagnostic of specific facial expression sounds, in both groups, but not of specific speech sounds or the gender of the two actors/speakers (Fig S12). In summary, the pattern of results observed in the right pSTS was similar to the pattern detected in the FFA in blind subjects. Such a correspondence was not observed between the FFA and other control ROIs.

**Figure 6.**
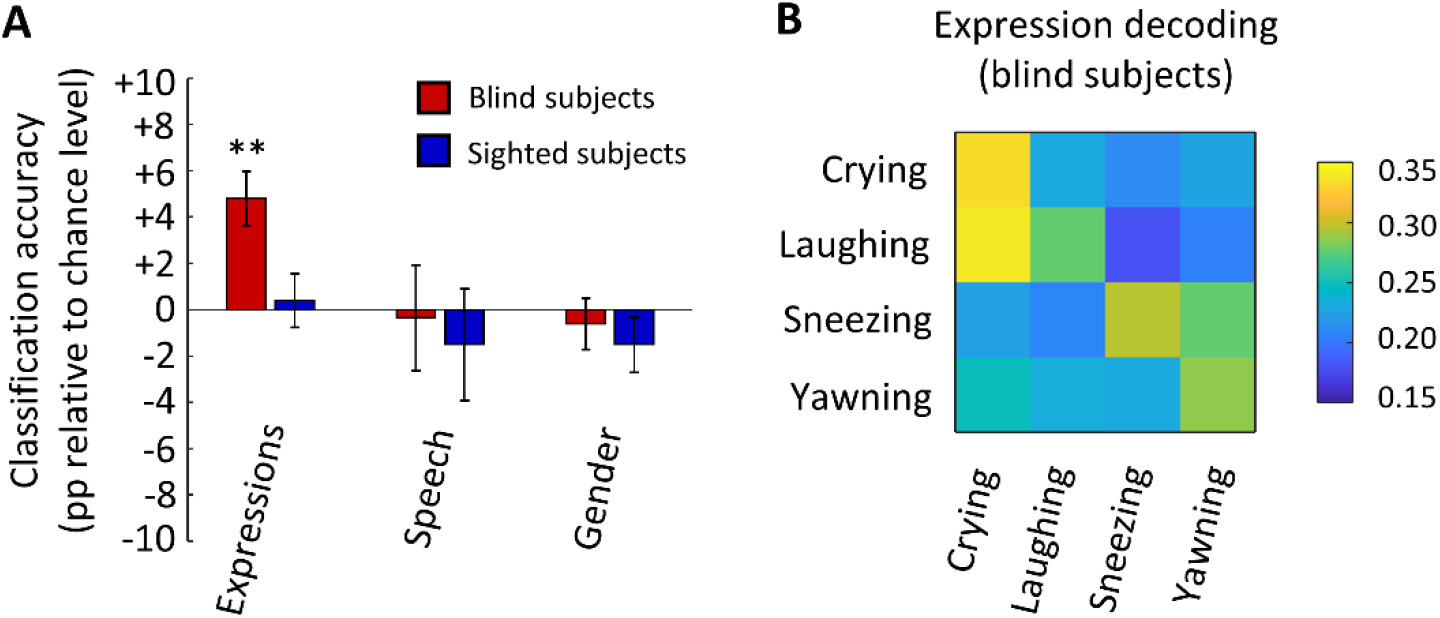
Activation patterns in the fusiform gyrus of blind subjects code differences between specific facial expressions presented through the auditory modality. (A) Mean accuracy of a support vector machine classifier distinguishing, based on the activation in the typical location of the FFA, between: four specific facial expressions (crying, laughing, sneezing and yawning), irrespectively of the actor; two specific speech sounds, irrespectively of the actor; and the actor (a man and a woman), irrespectively of the specific facial expression or the speech sound. The presented accuracy is adjusted for the chance level (25 % for facial expression classification; 50 % for speech sound and gender classification). The FFA was defined based on the Neurosynth meta-analysis for the term “face” (896 studies included; the resulting map thresholded at Z = 10; for the purpose of the classification analysis, voxels in both the right and the left hemisphere were included, see Methods). Testing against chance was performed in a permutation analysis. (B) A confusion matrix representing the classifier predictions for each facial expression sound in the blind subjects. Rows of the confusion matrix indicate the facial expression sound presented during the experiment and columns indicate classifier’s predictions for this sound. The classifier’s predictions are represented by color hues, with warmer colors for higher proportion of predictions and colder colors for lower proportion of predictions. Statistical thresholds: (A) ** p < 0.01, corrected for multiple comparisons across all comparisons performed using FDR. Error bars represent standard error of the mean.

#### RSA

Finally, we used RSA to directly investigate similarities between response patterns induced in the FFA by different sound categories. Based on our hypothesis, we created a simple theoretical model assuming that responses to both types of facial expression sounds are the most similar to each other (animate sounds with a clear mapping between a facial motor action and a face shape), somewhat similar to speech sounds (animate sounds for which a mapping between a facial motor action and a face shape is decodable only for the mouth region, and therefore is less salient), and the least similar to animal and object sounds (animate sounds with no clear mapping between a facial motor action and a face shape and inanimate sounds) (see Fig. 7A for the graphical illustration of the model). We observed a significant correlation between this theoretical model and FFA response patterns in the blind group (p_FDR_ = 0.012), but not in the sighted group (p_FDR_ = 0.223) (Fig. 7B).

**Figure 7.**
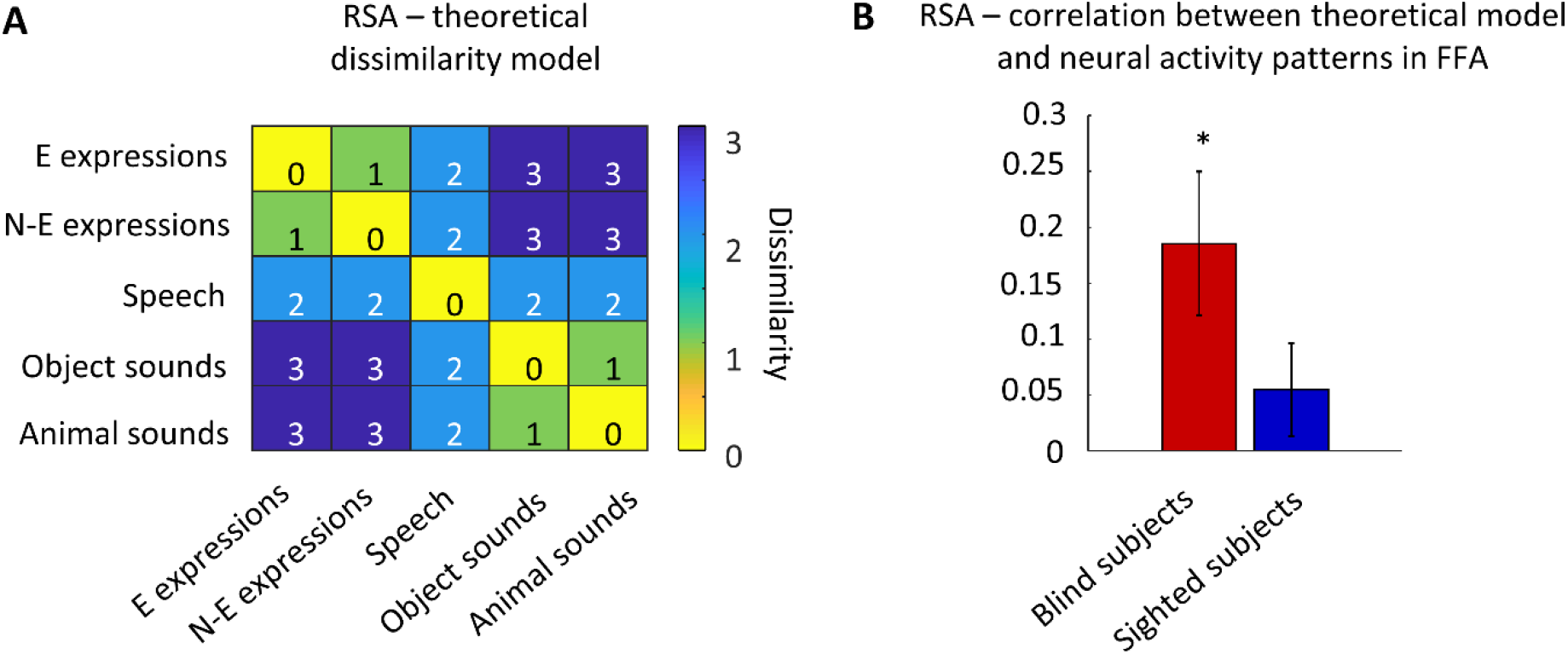
Similarity in activation patterns in the fusiform gyrus of blind subjects correlates with transparency of mapping between face shape and face action representations. (A) A theoretical model of FFA responses to sound categories used in the study. The categories with higher expected dissimilarity in responses (based on the study hypothesis) were assigned higher dissimilarity scores and are marked with colder color hues. E expressions - emotional expression sounds, N-E expressions – non-emotional expression sounds. (B) A representational similarity analysis (RSA) - correlation between the theoretical model and observed FFA responses. Testing against chance was performed in a permutation analysis. * p = 0.012, corrected for the two tests performed using FDR.

## Discussion

In this study, we found that sounds associated with facial expressions induce preferential response in the lateral fusiform gyrus - the typical anatomical location of the FFA - in congenitally blind individuals. This effect is independent of the emotional content (or lack thereof) of the facial expression sounds and was observed in comparisons to object sounds, animal sounds, and human speech. In contrast to facial expression sounds, speech sounds or animal sounds did not yield preferential activation, relative to object sounds, in the vOTC animate areas in the blind group. In the sighted subjects, no clear functional preference in the vOTC was observed for any of the animate sound categories. However, the multi-voxel pattern classification showed that the expected distinctions between facial expression sounds and other sound categories are, to some extent, represented in the lateral fusiform gyrus in both groups, with the representation in blind subjects being precise enough to differentiate between specific facial expressions. The RSA showed that the theoretical model of FFA responses, derived from our visual-motor mapping hypothesis, correlates with the observed responses in this area in the blind but not in the sighted subjects.

In the visual modality, the FFA responses are known to be the strongest for human faces. However, responses to many other animate categories, such as animals’ heads, whole animals, whole humans or human bodies without heads were reported to be still stronger than responses to inanimate objects (Kanwisher et al., 1999). In our study, the univariate analysis revealed a more dichotomous pattern of results, with both types of facial expression sounds activating the FFA in blind subjects more strongly than the other categories, and the activation for speech sounds and animal sounds being not significantly different from the activation for object sounds. This clear dissociation brings a new perspective to a recent debate concerning this area’s sensitivity to auditory and tactile stimulation. Several studies have already shown that the FFA can be preferentially activated by auditory and tactile stimuli related to the human face, compared to other types of information conveyed through these sensory modalities (blind subjects: van den Hurk et al., 2017; Murty et al., 2020; sighted subjects: Kitada et al., 2009). However, other studies investigating this issue reported null results, and a theoretical explanation of this difference has been lacking (blind and sighted subjects: Goyal et al., 2006; Kitada et al., 2013; Plaza et al., 2015; Dormal et al., 2018). Here, in an extension of the conjecture formulated by Bi and colleagues (Bi et al., 2016), we proposed that these conflicting findings reflect the fact that various face shape representations are differently associated with facial motor actions. This view predicts that dynamic face shape representations, which are clearly related to specific facial actions, are more available through non-visual modalities than other, more static face shape representations. Recent auditory studies have already hinted at this difference: null effect in the fusiform gyrus of blind and sighted subjects was reported in a study using vowels spoken in a neutral tone – a stimuli for which a mapping between the facial motor action and the resulting face shape is decodable only for the mouth region, and therefore is not particularly salient (Dormal et al., 2018); in contrast, a significant increase in the FFA activation in blind subjects was reported for sounds of facial actions (such as laughing, chewing, blowing a kiss or whistling; van den Hurk et al., 2017) and for verbal phrases conveying information about a speaker’s emotional state (Fairhall et al., 2017) – that is, dynamic stimuli for which a link between the facial motor action and the resulting face shape is clear and salient. In line with these results, our study shows a clear preference of the FFA in the blind group for facial expression sounds. This preference we observed independently of whether emotional content was present or not, which speaks against the recruitment of the FFA in blind individuals for emotional processing (e.g., Fairhall et al., 2017). These findings confirm our initial hypothesis and suggests that the transparency of mapping between visual and action representations modulates the auditory responsiveness of the FFA. Furthermore, our findings contribute to a growing body of evidence that visual experience is not necessary for the development of functional preference for face-related information in the human fusiform gyrus (van den Hurk et al., 2017; Murty et al., 2020) and, more broadly, for the development of a relatively typical functional organization in the OTC (e.g., Pietrini et al., 2004; Mahon et al., 2009; Wolbers et al., 2011; He et al., 2013, Peelen et al., 2013; Ricciardi et al., 2014; Wang et al., 2015).

On a more general level, our study contributes to the understanding of the nature of a perplexing phenomenon – that is, the sensory modality by object domain interaction in the OTC. A recent conjecture proposed that weaker auditory and tactile responsiveness of vOTC animate areas, relative to inanimate areas, can be explained by the fact that the mapping between visual and action representations for the animate domain is less transparent than for inanimate objects (Bi et al., 2016). In this study, we demonstrate the predictive value of this hypothesis by showing that effects characteristic of the inanimate domain can be also observed for specific subsets of animate entities with clear relationship between visual and action representations. Our findings suggest that the OTC organization is determined not only by local sensitivity to certain visual features, but also by metrics computed downstream in the brain, particularly in the action system (Mahon et al., 2007; Mahon & Caramazza, 2011).

An unexpected finding in our study is the clear difference in vOTC univariate selectivity to facial expression sounds across the congenitally blind and the sighted group. This seems to be inconsistent with the results from the inanimate domain, which are remarkably similar for these two groups (He et al., 2013; Peelen et al., 2013; Wang et al., 2015). The stronger selectivity to facial expression sounds in the blind group was observed despite the fact that, arguably, congenitally blind subjects cannot form as precise mental model of a human face as sighted subjects – thus, mental imagery cannot account for this result. One might take this between-group difference as evidence for functional repurposing (i.e., change in the type of representation computed) of this cortical territory, following the loss of sight (Bedny, 2017; Fairhall et al., 2017; Livingstone et al., 2019). However, postulating this kind of a change would require a plausible account of a stimulus dimension, different than the face shape and its relationship with face action representations, that is captured by this region in blind subjects in our study. Since the effect observed in the fusiform gyrus in the blind group seems to be independent from auditory, linguistic, social, and emotional content of sounds, it is unclear what this dimension might be. Based on these considerations, and on our observation of generally weaker response to sounds in the FFA of the sighted, we think that the adaptation of the FFA circuitry to the strength of incoming signals is a plausible explanation of the observed between-group difference. In sighted individuals, the FFA receives strong feedforward inputs from early visual cortices, which might make this region relatively insensitive to small differences between inputs from other brain systems. In congenitally blind individuals, lack of feedforward signals might lead to the adjustment of the FFA sensitivity – as a result, this area might be able to capture the subtle differences induced by auditory stimuli. Such an adjustment of sensitivity, following lack of sight, might be weaker for the inanimate representations, computed in other parts of the vOTC, as these representations are generally more available to information from other brain systems (see the discussion above). In agreement with our findings, the two previous studies that showed univariate activation in the fusiform gyrus of blind individuals, induced by auditory stimulation, failed to find similar, robust effects in this region in sighted individuals (Fairhall et al., 2017; van den Hurk et al., 2017). Overall, while the difference between the blind and the sighted subjects was not predicted in our study, it does not affect the theoretical implications of our results, described above – unless some form of functional repurposing in the fusiform gyrus in the blind group is assumed (e.g., Fairhall et al., 2017), for which, we argue, our study provides no convincing evidence.

In conclusion, we show that auditory responsiveness of the lateral fusiform gyrus (the typical anatomical location of the FFA) in congenitally blind individuals is modulated by the type of evoked face shape representation and its relationship with face action representations. Thus, OTC representation of both inanimate and animate stimuli can be reliably activated by auditory signals as long as the mapping between shape and associated action/motor computations is stable and transparent.

## Funding

This work was supported by a Polish Ministry of Science grant (DN/MOB/023/V/2017) and a Kosciuszko Foundation fellowship to Ł.B., the National Natural Science Foundation of China (31925020, 31671128 to Y.B.), Changjiang Scholar Professorship Award (T2016031 to Y.B.), the 111 Project (BP0719032 to Y.B.), and the Fundamental Research Funds for the Central Universities.

## Competing interests

The authors declare that no competing interests exists.

## Notes

### Competing Interest Statement

The authors have declared no competing interest.

## References

Andrews TJ, Ewbank MP. 2004. Distinct representations for facial identity and changeable aspects of faces in the human temporal lobe. Neuroimage. 23: 905–913.

Allison T, Puce A, McCarthy G. 2000. Social perception from visual cues: role of the STS region. Trends Cogn Sci. 4: 267–278

Bedny M. 2017. Evidence from Blindness for a Cognitively Pluripotent Cortex. Trends Cogn Sci. 21:637–648.

Benjamini Y, Hochberg Y. 1995. Controlling the False Discovery Rate: A Practical and Powerful Approach to Multiple Testing. J R Stat Soc Ser B. 57:289–300.

Bi Y, Wang X, Caramazza A. 2016. Object Domain and Modality in the Ventral Visual Pathway. Trends Cogn Sci. 20:282–290.

Bischoff-Grethe A, Ozyurt IB, Busa E, Quinn BT, Fennema-Notestine C, Clark CP, Morris S, Bondi MW, Jernigan TL, Dale AM, Brown GG, Fischl B. 2007. A technique for the deidentification of structural brain MR images. Hum Brain Mapp. 28:892–903.

Chang CC, Lin CJ. 2011. LIBSVM: A Library for support vector machines. ACM Trans Intell Syst Technol. 2:27.

Cousineau D. 2005. Confidence intervals in within-subject designs: A simpler solution to Loftus and Masson’s method. Tutor Quant Methods Psychol. 1: 42–45.

Dormal G, Pelland M, Rezk M, Yakobov E, Lepore F, Collignon O. 2018. Functional Preference for Object Sounds and Voices in the Brain of Early Blind and Sighted Individuals. J Cogn Neurosci. 30:86–106.

Epstein R, Kanwisher N. 1998. A cortical representation of the local visual environment. Nature. 392: 598–601.

Fairhall SL, Anzellotti S, Ubaldi S, Caramazza A. 2014. Person- and Place-Selective Neural Substrates for Entity-Specific Semantic Access. Cereb Cortex. 24:1687–1696.

Fairhall SL, Porter KB, Bellucci C, Mazzetti M, Cipolli C, Gobbini MI. 2017. Plastic reorganization of neural systems for perception of others in the congenitally blind. Neuroimage. 158:126–135.

Friston KJ, Holmes AP, Poline JB, Grasby PJ, Williams SC, Frackowiak RS, Turner R. 1995. Analysis of fMRI time-series revisited. Neuroimage. 2:45–53.

Gardumi A, Ivanov D, Hausfeld L, Valente G, Formisano E, Uludağ K. 2016. The effect of spatial resolution on decoding accuracy in fMRI multivariate pattern analysis. Neuroimage. 132:32–42.

Gauthier I, Tarr MJ, Moylan J, Skudlarski P, Gore JC, Anderson AW. 2000. The fusiform “face area” is part of a network that processes faces at the individual level. J Cogn Neurosci. 12:495–504.

Goyal MS, Hansen PJ, Blakemore CB. 2006. Tactile perception recruits functionally related visual areas in the late-blind. Neuroreport. 17:1381–1384.

He C, Peelen M V, Han Z, Lin N, Caramazza A, Bi Y. 2013. Selectivity for large nonmanipulable objects in scene-selective visual cortex does not require visual experience. Neuroimage. 79:1–9.

Ishai A, Ungerleider LG, Haxby JV. 2000. Distributed neural systems for the generation of visual images. Neuron. 28:979–990.

Kanwisher N, McDermott J, Chun MM. 1997. The fusiform face area: a module in human extrastriate cortex specialized for face perception. J Neurosci. 17:4302–4311.

Kanwisher N, Stanley D, Harris A. 1999. The fusiform face area is selective for faces not animals. Neuroreport. 10:183–187.

Kanwisher N, Yovel G. 2006. The fusiform face area: a cortical region specialized for the perception of faces. Philos Trans R Soc B Biol Sci. 361:2109–2128.

Kitada R, Johnsrude IS, Kochiyama T, Lederman SJ. 2009. Functional specialization and convergence in the occipito-temporal cortex supporting haptic and visual identification of human faces and body parts: An fMRI study. J Cogn Neurosci. 21:2027–2045.

Kitada R., Okamoto Y., Sasaki AT., Kochiyama, T., Miyahara, M., Lederman, S. J., Sadato, N. 2013. Early visual experience and the recognition of basic facial expressions: involvement of the middle temporal and inferior frontal gyri during haptic identification by the early blind. Front Hum Neurosci. 7: 7.

Livingstone MS, Arcaro MJ, Schade PF. 2019. Cortex Is Cortex: Ubiquitous Principles Drive Face-Domain Development. Trends Cogn Sci. 23:3–4.

Magri C, Konkle T, Caramazza A. 2020. The contribution of object size, manipulability, and stability on neural responses to inanimate objects. bioRxiv.

Mahon BZ, Milleville SC, Negri GA, Rumiati RI, Caramazza A, Martin A. 2007. Action-related properties shape object representations in the ventral stream. Neuron. 55: 507–520.

Mahon BZ, Anzellotti S, Schwarzbach J, Zampini M, Caramazza A. 2009. Category-specific organization in the human brain does not require visual experience. Neuron. 63:397–405.

Mahon BZ, Caramazza A. 2011. What drives the organization of object knowledge in the brain?. Trends Cogn Sci. 15: 97–103.

Murty NAR, Teng S, Beeler D, Mynick A, Oliva A, Kanwisher NG. 2020. Visual Experience is not Necessary for the Development of Face Selectivity in the Lateral Fusiform Gyrus. Proc Natl Acad Sci USA. 117: 23011–23020.

O’Craven KM, Kanwisher N. 2000. Mental imagery of faces and places activates corresponding stimulus-specific brain regions. J Cogn Neurosci. 12:1013–1023.

Oosterhof NN, Connolly AC, Haxby J V. 2016. CoSMoMVPA: Multi-Modal Multivariate Pattern Analysis of Neuroimaging Data in Matlab/GNU Octave. Front Neuroinform. 10.

Peelen MV, Bracci S, Lu X, He C, Caramazza A, Bi Y. 2013. Tool Selectivity in Left Occipitotemporal Cortex Develops without Vision. J Cogn Neurosci. 25:1225–1234.

Peirce JW. 2007. PsychoPy-Psychophysics software in Python. J Neurosci Methods. 162:8–13.

Pietrini P, Furey ML, Ricciardi E, Gobbini MI, Wu WHC, Cohen L,… Haxby JV. 2004. Beyond sensory images: Object-based representation in the human ventral pathway. Proc Natl Acad Sci USA. 101: 5658–5663.

Pitcher D, Duchaine B, Walsh V. 2014. Combined TMS and fMRI reveal dissociable cortical pathways for dynamic and static face perception. Curr Biol. 24: 2066–2070.

Plaza P, Renier L, De Volder A, Rauschecker J. 2015. Seeing faces with your ears activates the left fusiform face area, especially when you’re blind. J Vis. 15: 197–197.

Ricciardi E, Bonino D, Pellegrini S, Pietrini P. 2014. Mind the blind brain to understand the sighted one! Is there a supramodal cortical functional architecture?. Neurosc Biobehav Rev. 41: 64–77.

Smith SM, Nichols TE. 2009. Threshold-free cluster enhancement: Addressing problems of smoothing, threshold dependence and localisation in cluster inference. Neuroimage. 44:83–98.

van den Hurk J, Van Baelen M, Op de Beeck HP. 2017. Development of visual category selectivity in ventral visual cortex does not require visual experience. Proc Natl Acad Sci USA. 114:E4501–E4510.

Wang X, Peelen M V, Han Z, He C, Caramazza A, Bi Y. 2015. How Visual Is the Visual Cortex? Comparing Connectional and Functional Fingerprints between Congenitally Blind and Sighted Individuals. J Neurosci. 35:12545–12559.

Whitfield-Gabrieli S, Nieto-Castanon A. 2012. Conn: A Functional Connectivity Toolbox for Correlated and Anticorrelated Brain Networks. Brain Connect. 2:125–141.

Wolbers T, Klatzky RL, Loomis JM, Wutte MG, Giudice NA. 2011. Modality-Independent Coding of Spatial Layout in the Human Brain. Curr Biol. 21:984–989.

Yarkoni T, Poldrack RA, Nichols TE, Van Essen DC, Wager TD. 2011. Large-scale automated synthesis of human functional neuroimaging data. Nat Methods. 8:665–670.

