## Supplementary Figures and Tables for "Preference for animate domain sounds in the fusiform gyrus of blind individuals is modulated by shape-action mapping"

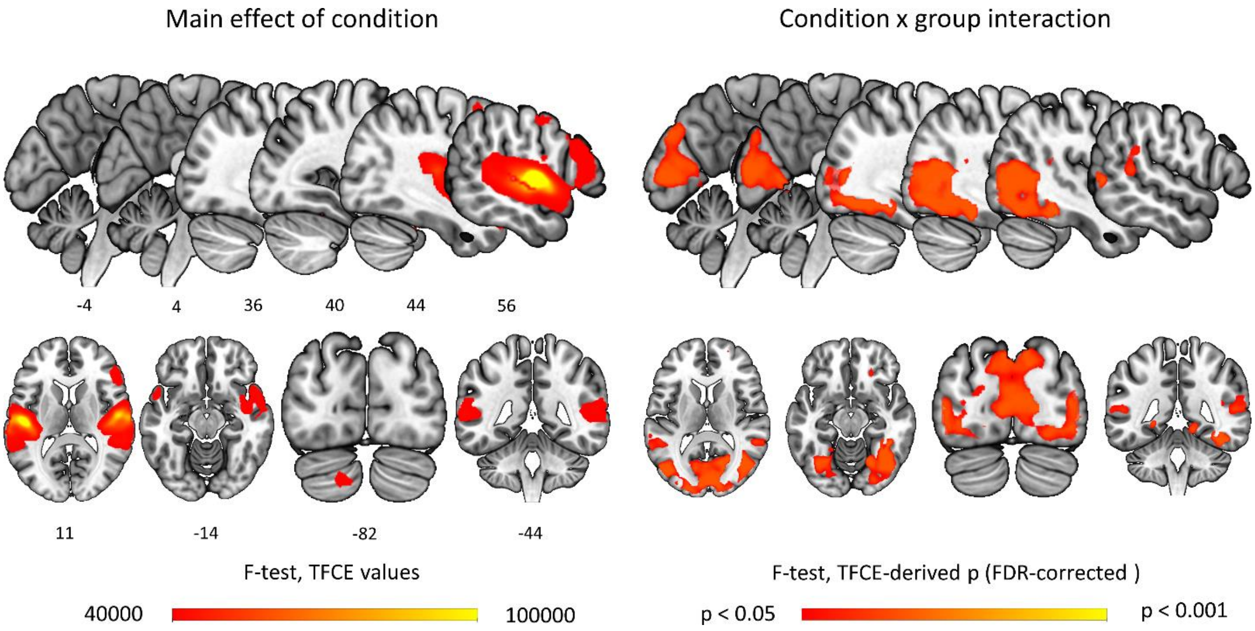


**Figure S1. Results of omnibus F-tests.** Testing for the main effect of group did not yield significant results. Statistical thresholds: Initially, all analysis parameters were identical to those applied to the data reported in the main text (i.e., statistical threshold was set at false-discovery-rate corrected $p < 0.05$, derived based on calculation of TFCE values and permutation testing; see Figure 1 for details). In the case of the testing for the main effect of condition, this yielded significant effects in wide network of regions, including frontoparietal regions, the temporal lobe, the OTC, and the early visual cortex (see Table S2). To achieve a higher degree of spatial specificity, the statistical threshold for this analysis was increased to arbitrarily chosen TFCE value (i.e., 40000).

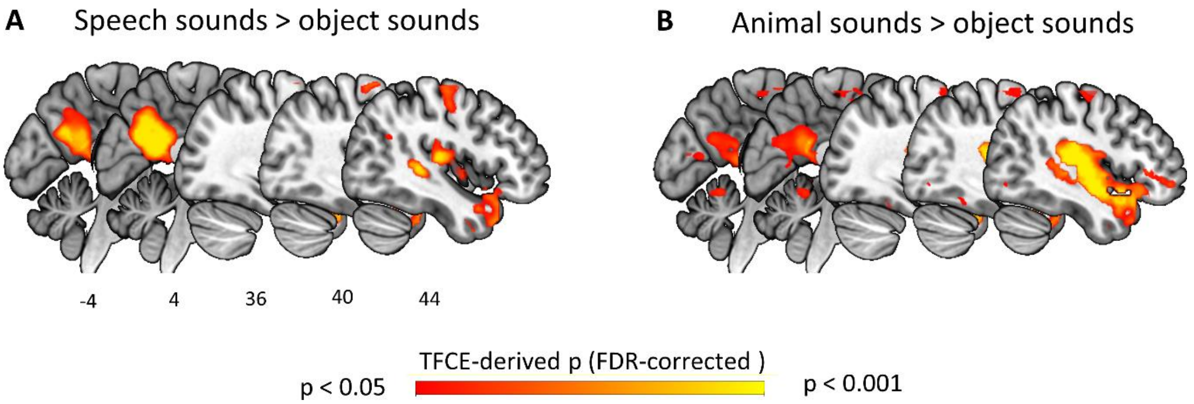


**Figure S2. Preferential activation induced by speech sounds (A) or animal sounds (B), relative to object sounds, in blind subjects.** All analysis parameters were identical to those applied to the data reported in the main text (see Figure 2 for details).

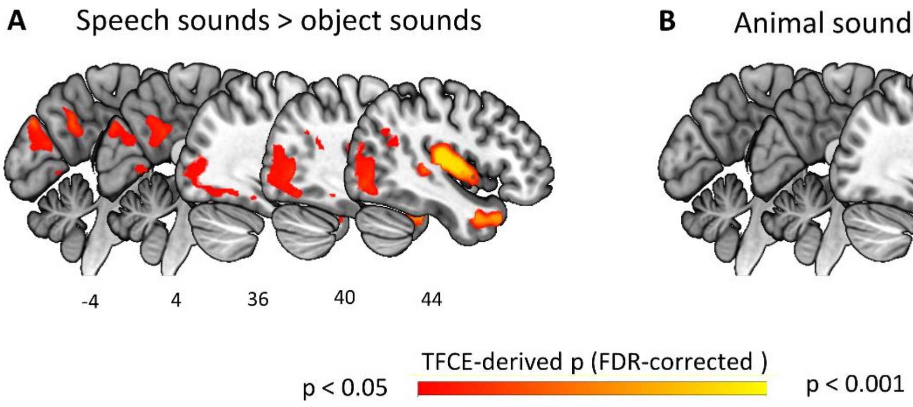


**Figure S3. Preferential activation induced by speech sounds (A) or animal sounds (B), relative to object sounds, in sighted subjects.** All analysis parameters were identical to those applied to the data reported in the main text (see Figure 2 for details).

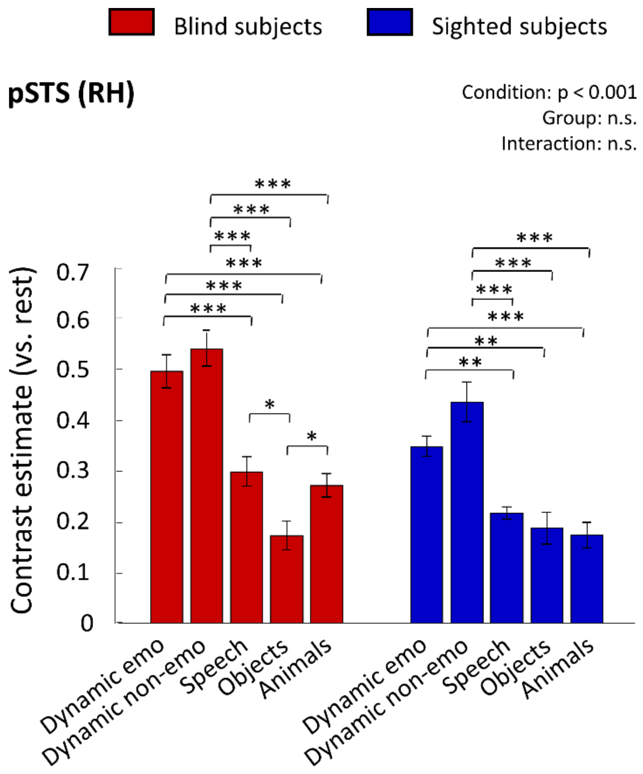


**Figure S4. Region-of-interest (ROI) analysis of the univariate activation patterns in the right posterior superior temporal sulcus (right pSTS).** Mean contrast estimates, calculated relative to rest periods, are presented for each condition and group for the right pSTS. The ROI was defined based on the Neurosynth metanalysis of studies including the words "facial expressions" (250 studies included; map thresholded at Z = 4.8; ROI size = 233 functional voxels). Condition x Group ANOVA showed a significant effect of condition ($p < 0.001$; all other ANOVA effects were not significant). ** $p < 0.01$, *** $p < 0.001$, corrected for multiple comparisons using the false discovery rate. Error bars represent standard error of the mean, adjusted to adequately represent within-subject variability across conditions, in each group, using a method proposed by Cousineau (2005).

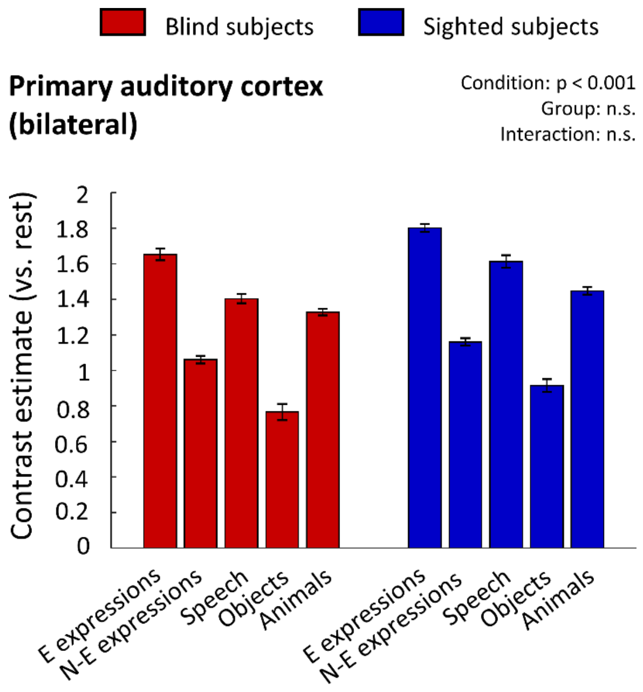


**Figure S5. Region-of-interest (ROI) analysis of the univariate activation patterns in the primary auditory cortex (A1).** Mean contrast estimates, calculated relative to rest periods, are presented for each condition and group for the primary auditory cortex. The ROI was defined bilaterally, based on the Neurosynth metanalysis of studies including the words "primary auditory" (114 studies included; map thresholded at Z = 10; ROI size = 672 functional voxels). Condition x Group ANOVA showed a significant effect of condition ($p < 0.001$; all other ANOVA effects were not significant). Responses to all conditions are significantly different from each other (all $p < 0.001$). Error bars represent standard error of the mean, adjusted to adequately represent within-subject variability across conditions, in each group, using a method proposed by Cousineau (2005).

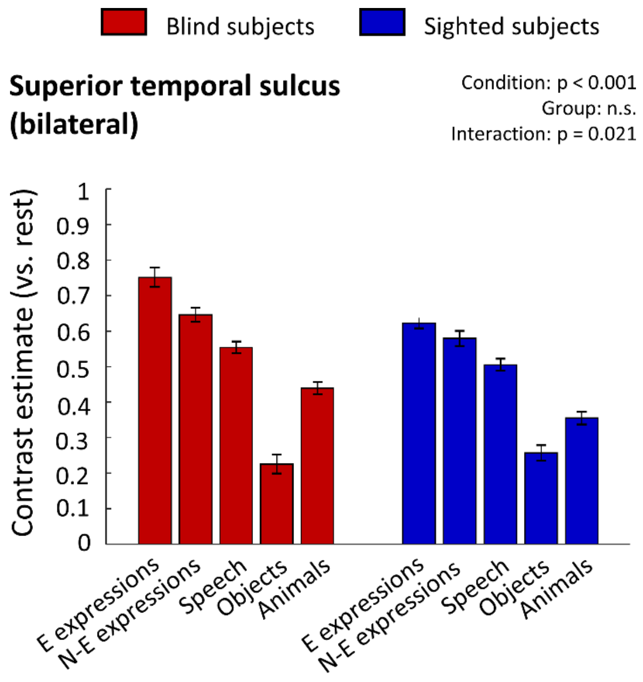


**Figure S6. Region-of-interest (ROI) analysis of the univariate activation patterns in the superior temporal sulcus (STS).** Mean contrast estimates, calculated relative to rest periods, are presented for each condition and group for the primary auditory cortex. The ROI was defined bilaterally, based on the Neurosynth metanalysis of studies including the abbreviation "STS" (203 studies included; map thresholded at $Z = 7$; ROI size = 2234 functional voxels). Condition x Group ANOVA showed a significant effect of condition ($p < 0.001$) and a significant condition x group interaction ($p = 0.021$). Responses to all conditions are significantly different from each other (all false-discovery-rate-corrected $p < 0.01$), except from responses to emotional and non-emotional expression sounds in the sighted subjects ($p > 0.15$). Error bars represent standard error of the mean, adjusted to adequately represent within-subject variability across conditions, in each group, using a method proposed by Cousineau (2005).

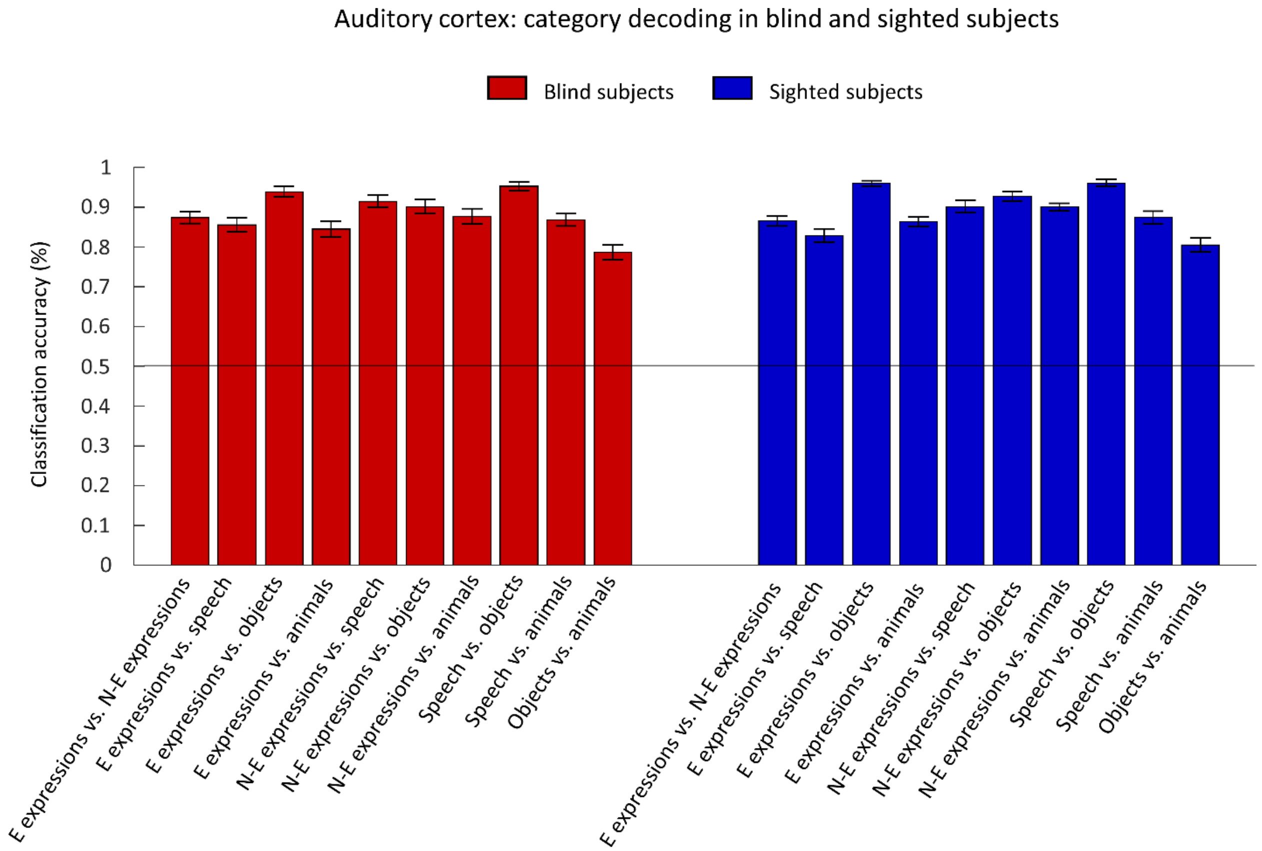


**Figure S7. Classification of sound categories based on the activation of the auditory cortex.** The auditory cortex ROI was defined as the A1 ROI and the STS ROI combined (see Fig. S5 and S6). All analysis parameters were identical to those described in Fig. 5. All sound category pairs were distinguished with high accuracy (p = 0.001; chance level is marked with a black line).

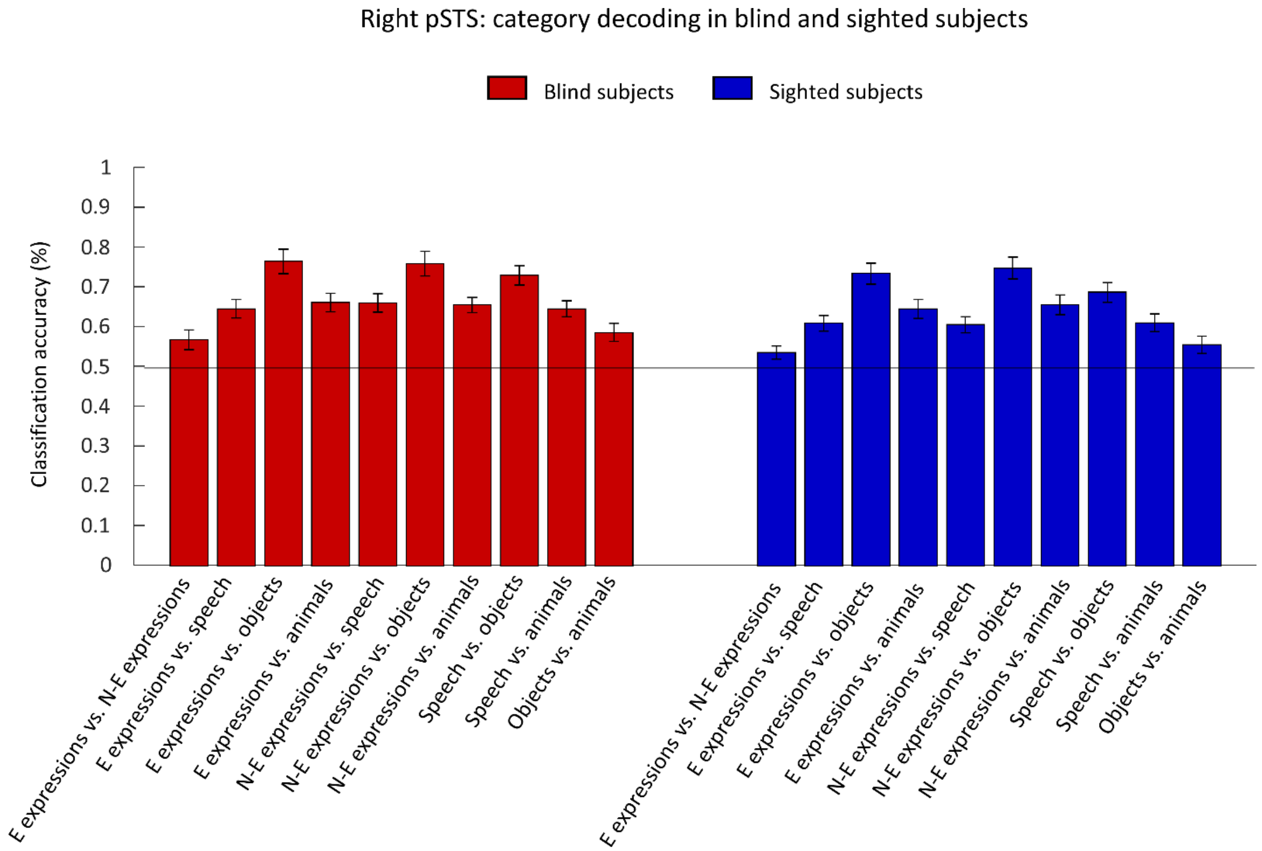


**Figure S8. Classification of sound categories based on the activation of the right pSTS.** The right pSTS ROI was defined based on the Neurosynth analysis of studies using a term "facial expressions" (250 studies included; map thresholded at $Z = 2$, size = 440 functional voxels). All analysis parameters were identical to those described in Fig. 5. All sound category pairs were successfully distinguished ($p < 0.05$, false-discovery-rate for multiple comparisons across all tests performed within each group). Chance level is marked with a black line.

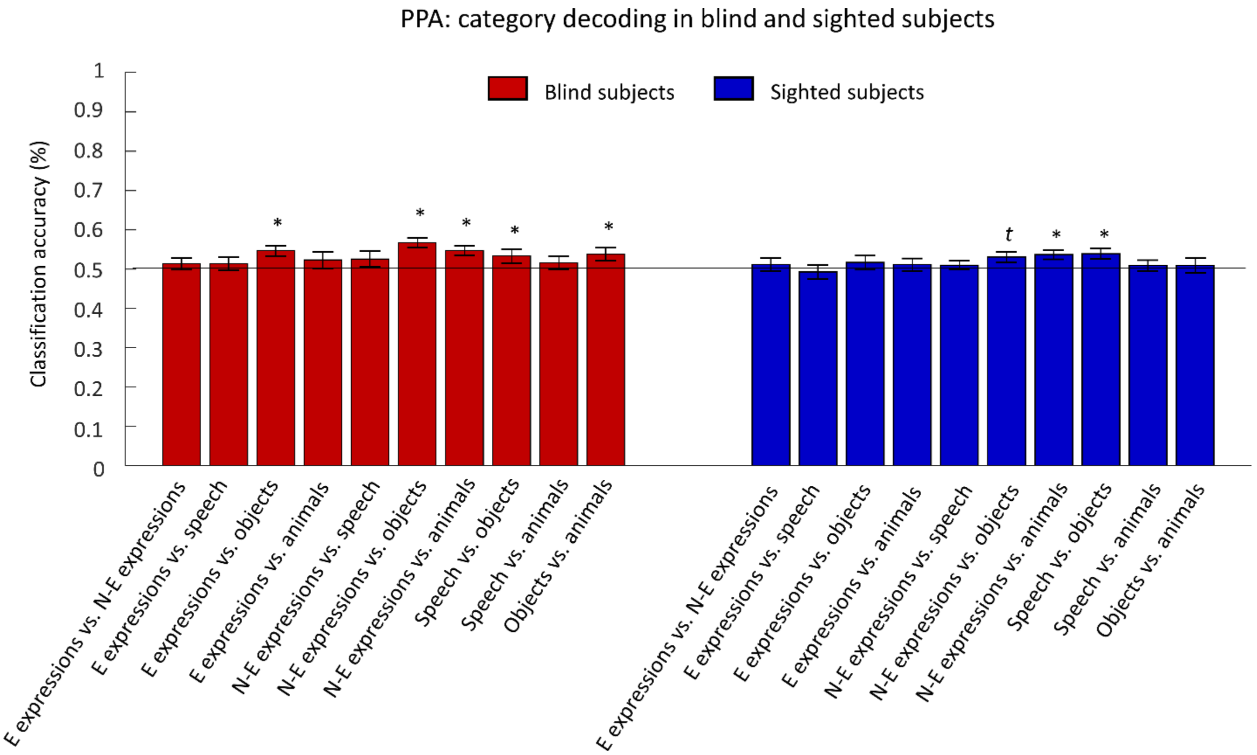


**Figure S9. Classification of sound categories based on the activation of the parahippocampal place area (PPA).** The PPA ROI was defined bilaterally based on the Neurosynth analysis of studies using a term "place" (189 studies included; map thresholded at $Z = 3$, size = 790 functional voxels). All analysis parameters were identical to those described in Fig. 5. * $p < 0.05$, $t = 0.063$, false-discovery-rate for multiple comparisons across all tests performed within each group.

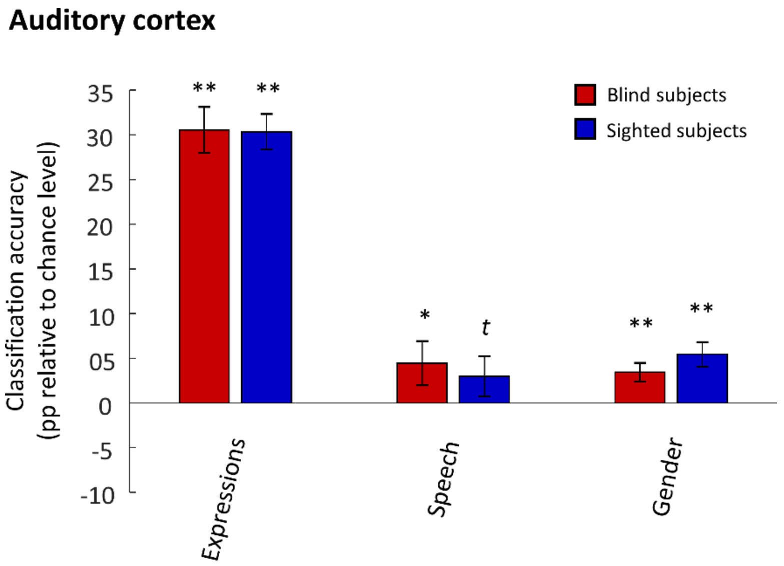


**Figure S10. Classification based on the activation of the auditory cortex.** Auditory cortex was defined as the A1 and the STS ROIs combined (see Fig. S5 and S6). All analysis parameters were identical to those described in Fig. 6. * $p < 0.05$, ** $p < 0.01$, $t = 0.085$, FDR-corrected for multiple comparisons.

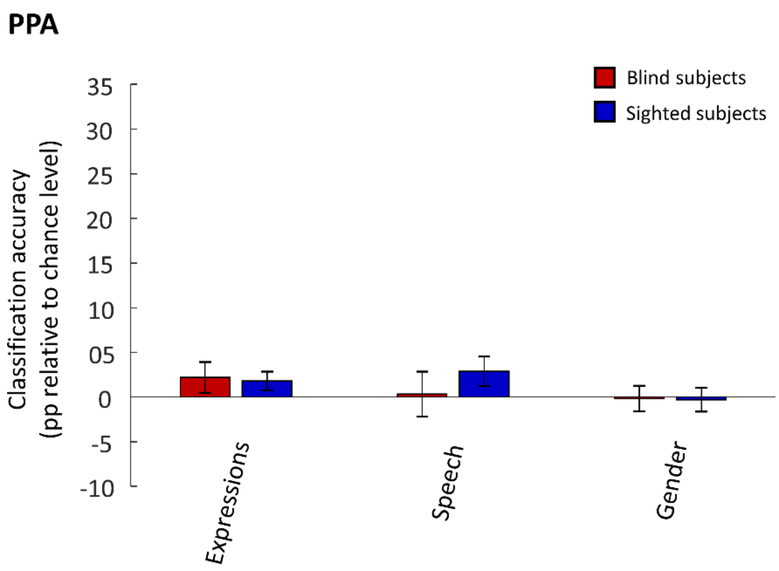


**Figure S11. Classification based on the activation of the PPA.** The PPA ROI was defined bilaterally based on the Neurosynth analysis of studies using a word "place" (189 studies included; map thresholded at Z = 3, size = 790 functional voxels). All analysis parameters were identical to those described in Fig. 6. All false-discovery-rate-corrected $p > 0.18$.

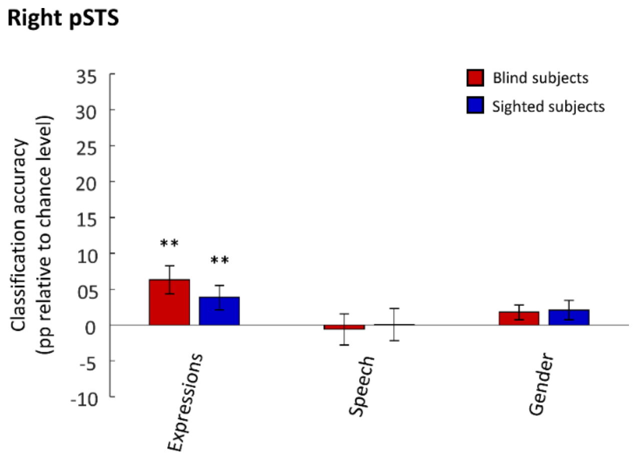


**Figure S12. Classification based on the activation of right pSTS.** The right pSTS ROI was defined based on the Neurosynth analysis of studies using a term "facial expressions" (250 studies included; map thresholded at Z = 2, size = 440 functional voxels). All analysis parameters were identical to those described in Fig. 6. ** $p < 0.01$, FDR-corrected for multiple comparisons.

**Table S1**. Results of the behavioral rating of stimuli, performed, on the independent group of subjects, before the actual fMRI study. Valence was rated on a 7-point Likert scale (1 - very negative; 7 - very positive). Sound recognizability was tested at the level of sound types ("is this sound made by a human, an animal or an artificial object?") and at the level of specific sound exemplars ("name a human action performed or an animal making this sound or an object making this sound"). Additionally, in the case of human sounds, subjects were asked to categorize emotional state of a person making a given sound ("is this person sad, happy, angry, scared or at no particular emotional state? Choose one answer that best describes his/her emotional state"). The table presents means and standard deviations.

| Sound type | Valence: rater | Valence: person/animal making the sound | Recognizability: category (human/animal/object) | Recognizability: specific action/specific animal/specific object | Recognizability: name the emotional state of person making the sound (sadness/happiness/anger/fear/neutral) |
|---|---|---|---|---|---|
| Emotional expression sounds: crying | 1.82 (0.11) | 1.69 (0.12) | 100% | 97% (2%) | 95% (3%) |
| Emotional expression sounds: laughing | 5.38 (0.23) | 5.92 (0.23) | 100% | 100% | 98 (4%) |
| Non-emotional expression sounds | 3.24 (0.13) | 3.25 (0.18) | 100% | 97% (4%) | 88% (4%) |
| Speech sounds | 4.06 (0.11) | 3.72 (0.12) | 100% | 100% | 88% (5%) |
| Object sounds | 3.91 (0.89) | NA | 100% | 92% (9%) | NA |
| Animal sounds | 3.97 (1.06) | 3.99 (1.02) | 100% | 97% (5%) | NA |

**Table S2**. F-test, the main effect of sound category. Only local maxima that are more than 24 mm apart are reported. TFCE - threshold free cluster enhancement; FDR - false discovery rate; MNI - Montreal Neurological Institute; Anatomical localization of MNI coordinates was performed using Anatomy SPM toolbox (v.2.2c).

| Region | TFCE value | p(FDR-corrected) | x | y | z (MNI space) |
|---|---|---|---|---|---|
| R Heschls Gyrus | 28372772.00 | 0.000 | 54 | -10 | 6 |
| L Superior Temporal Gyrus | 23395876.00 | 0.000 | -54 | -16 | 6 |
| R Superior Temporal Gyrus | 813906.38 | 0.000 | 50 | -34 | 4 |
| L Middle Temporal Gyrus | 291605.44 | 0.000 | -62 | -44 | 8 |
| R SupraMarginal Gyrus | 107848.05 | 0.000 | 64 | -38 | 24 |
| R IFG p. Triangularis | 79323.34 | 0.000 | 46 | 20 | 24 |
| R Temporal Pole | 75634.85 | 0.000 | 50 | 18 | -16 |
| R Precentral Gyrus | 65493.70 | 0.000 | 46 | 4 | 44 |
| L Precentral Gyrus | 45259.79 | 0.000 | -60 | 0 | 24 |
| L Cerebelum Crus 2 | 43363.66 | 0.000 | -12 | -80 | -42 |

| | | | | | |
|---|---|---|---|---|---|
| R IFG p. Triangularis | 43192.61 | 0.000 | 52 | 36 | 6 |
| R Medial Temporal Pole | 33486.96 | 0.000 | 30 | 18 | -34 |
| Undefined | 32493.03 | 0.000 | 30 | 26 | 2 |
| L Precentral Gyrus | 26022.84 | 0.000 | -50 | -4 | 50 |
| Undefined | 23779.52 | 0.000 | -34 | 2 | -20 |
| L Postcentral Gyrus | 19958.87 | 0.000 | -34 | -36 | 42 |
| R Middle Temporal Gyrus | 17421.52 | 0.000 | 66 | -10 | -20 |
| R Middle Temporal Gyrus | 17421.52 | 0.000 | 54 | -58 | 0 |
| R Precentral Gyrus | 17421.52 | 0.000 | 48 | -16 | 60 |
| L IFG p. Triangularis | 16749.07 | 0.000 | -40 | 24 | -2 |
| L SupraMarginal Gyrus | 16233.23 | 0.000 | -60 | -46 | 32 |
| L IFG p. Opercularis | 15001.42 | 0.000 | -36 | 10 | 28 |
| L Posterior-Medial Frontal | 14451.98 | 0.000 | -4 | -2 | 64 |
| L Postcentral Gyrus | 11423.24 | 0.000 | -50 | -22 | 30 |
| L Putamen | 10901.62 | 0.000 | -22 | 8 | 0 |
| L Cerebelum VIII | 10853.63 | 0.000 | -30 | -64 | -48 |
| L Middle Frontal Gyrus | 10418.26 | 0.000 | -32 | 50 | 26 |
| L IFG p. Triangularis | 10027.71 | 0.000 | -50 | 34 | 18 |
| L Superior Frontal Gyrus | 9611.63 | 0.000 | -18 | 64 | 10 |
| L Inferior Temporal Gyrus | 9360.13 | 0.000 | -48 | -62 | -6 |
| R Angular Gyrus | 9292.43 | 0.000 | 50 | -64 | 28 |
| R Mid Orbital Gyrus | 9280.61 | 0.000 | 4 | 50 | -12 |
| L Middle Orbital Gyrus | 9066.33 | 0.000 | -32 | 46 | -12 |
| L Middle Occipital Gyrus | 8802.66 | 0.000 | -30 | -76 | 38 |
| L Inferior Temporal Gyrus | 8665.01 | 0.000 | -62 | -8 | -26 |
| L Hippocampus | 8665.01 | 0.000 | -26 | -22 | -14 |
| L Angular Gyrus | 8665.01 | 0.000 | -50 | -68 | 26 |
| L Superior Medial Gyrus | 8665.01 | 0.000 | -2 | 58 | 28 |
| L Middle Frontal Gyrus | 8665.01 | 0.000 | -30 | 30 | 40 |
| L Precuneus | 8665.01 | 0.000 | -10 | -74 | 52 |
| L Superior Parietal Lobule | 8665.01 | 0.000 | -30 | -60 | 56 |
| L Postcentral Gyrus | 8665.01 | 0.000 | -48 | -28 | 60 |
| Undefined | 8187.77 | 0.000 | -22 | -12 | 50 |
| R Precuneus | 8077.24 | 0.000 | 6 | -56 | 22 |
| R Fusiform Gyrus | 7929.89 | 0.000 | 44 | -46 | -24 |
| R Hippocampus | 7462.22 | 0.000 | 20 | -8 | -14 |
| R Cerebelum VII | 6854.09 | 0.000 | 14 | -76 | -42 |
| L MCC | 5689.00 | 0.000 | -4 | 8 | 38 |
| R Posterior-Medial Frontal | 5531.81 | 0.000 | 8 | 24 | 52 |
| R Inferior Occipital Gyrus | 4977.23 | 0.001 | 42 | -78 | -10 |
| R Superior Frontal Gyrus | 4587.68 | 0.000 | 22 | 62 | 2 |
| L MCC | 4554.14 | 0.000 | -12 | -48 | 36 |
| R Precuneus | 4554.14 | 0.000 | 8 | -56 | 48 |
| Cerebellar Vermis 6 | 4436.82 | 0.000 | -2 | -76 | -18 |
| L Calcarine Gyrus | 4250.49 | 0.000 | 0 | -90 | 6 |

| | | | | | |
|---|---|---|---|---|---|
| L Middle Occipital Gyrus | 3993.47 | 0.000 | -42 | -86 | -2 |
| R Superior Occipital Gyrus | 3861.14 | 0.000 | 22 | -98 | 16 |
| L Cerebelum VI | 3674.22 | 0.001 | -34 | -70 | -24 |
| L Thalamus | 3593.06 | 0.000 | -10 | -20 | 6 |
| R Postcentral Gyrus | 3535.48 | 0.000 | 42 | -28 | 40 |
| R Middle Frontal Gyrus | 3517.74 | 0.000 | 32 | 38 | 42 |
| R Angular Gyrus | 3459.93 | 0.000 | 34 | -54 | 44 |
| R Putamen | 3345.22 | 0.000 | 26 | 2 | 8 |
| Undefined | 3250.06 | 0.000 | 8 | -30 | -12 |
| R Thal: Temporal | 3235.35 | 0.000 | 16 | -20 | 20 |
| L Olfactory cortex | 3213.68 | 0.000 | -2 | 24 | -4 |
| R ACC | 3181.59 | 0.000 | 4 | 22 | 20 |
| L Middle Occipital Gyrus | 3157.78 | 0.000 | -22 | -98 | 16 |
| Undefined | 3148.55 | 0.000 | 24 | 44 | 18 |
| R Posterior-Medial Frontal | 3095.07 | 0.000 | 8 | -24 | 62 |
| R Inferior Temporal Gyrus | 3051.81 | 0.000 | 48 | 2 | -36 |
| L Medial Temporal Pole | 3051.81 | 0.000 | -32 | 20 | -36 |
| Cerebellar Vermis 9 | 3051.81 | 0.000 | 0 | -58 | -34 |
| R Cerebelum Crus 1 | 3051.81 | 0.001 | 36 | -68 | -32 |
| L Cerebelum Crus 1 | 3051.81 | 0.000 | -40 | -48 | -32 |
| R Cerebelum IV-V | 3051.81 | 0.000 | 16 | -54 | -16 |
| L Middle Temporal Gyrus | 3051.81 | 0.000 | -52 | -28 | -16 |
| Undefined | 3051.81 | 0.000 | -4 | -4 | -16 |
| L Lingual Gyrus | 3051.81 | 0.000 | -16 | -58 | -10 |
| R Calcarine Gyrus | 3051.81 | 0.001 | 16 | -70 | 4 |
| L Calcarine Gyrus | 3051.81 | 0.001 | -12 | -70 | 14 |
| L Insula Lobe | 3051.81 | 0.000 | -32 | -14 | 16 |
| R Cuneus | 3051.81 | 0.001 | 16 | -78 | 28 |
| R Postcentral Gyrus | 3051.81 | 0.000 | 66 | -10 | 28 |
| R MCC | 3051.81 | 0.001 | 2 | -28 | 38 |
| R Angular Gyrus | 3051.81 | 0.001 | 36 | -78 | 42 |
| Undefined | 3051.81 | 0.000 | 26 | -10 | 46 |
| L Superior Medial Gyrus | 3051.81 | 0.000 | -8 | 42 | 46 |
| L Superior Frontal Gyrus | 3051.81 | 0.001 | -22 | 12 | 54 |
| R Superior Frontal Gyrus | 3051.81 | 0.000 | 28 | 12 | 58 |
| L Fusiform Gyrus | 3321.98 | 0.000 | -24 | -36 | -18 |
| L Inferior Temporal Gyrus | 3195.29 | 0.000 | -38 | -16 | -30 |
| R Fusiform Gyrus | 3051.81 | 0.000 | 30 | -68 | -4 |
| R Middle Occipital Gyrus | 3051.81 | 0.001 | 24 | -88 | 6 |

**Table S3**. F-test, the sound category by group interaction effect. Only local maxima that are more than 24 mm apart are reported. TFCE - threshold free cluster enhancement; FDR - false discovery rate; MNI - Montreal Neurological Institute; Anatomical localization of MNI coordinates was performed using Anatomy SPM toolbox (v.2.2c).

| Region | TFCE value | p(FDR-corrected) | x | y | z (MNI space) |
|---|---|---|---|---|---|
| R Middle Temporal Gyrus | 5809.65 | 0.013 | 46 | -70 | 4 |
| L Middle Occipital Gyrus | 5668.57 | 0.013 | -46 | -72 | 4 |
| R Lingual Gyrus | 5254.18 | 0.014 | 14 | -64 | 4 |
| L Calcarine Gyrus | 5052.81 | 0.014 | -6 | -90 | 6 |
| R Fusiform Gyrus | 4122.40 | 0.013 | 42 | -42 | -20 |
| L Lingual Gyrus | 3763.81 | 0.014 | -16 | -58 | -4 |
| L Cuneus | 3452.61 | 0.015 | -10 | -90 | 34 |
| R Lingual Gyrus | 3105.96 | 0.015 | 26 | -84 | -10 |
| R Cerebelum IV-V | 2126.63 | 0.015 | 14 | -44 | -10 |
| R Superior Occipital Gyrus | 2112.30 | 0.023 | 22 | -92 | 26 |
| L Fusiform Gyrus | 1860.15 | 0.037 | -40 | -62 | -20 |
| L Lingual Gyrus | 1743.20 | 0.039 | -22 | -80 | -14 |
| L Middle Occipital Gyrus | 1111.04 | 0.034 | -34 | -94 | 6 |
| L Middle Temporal Gyrus | 2493.96 | 0.013 | -62 | -26 | 0 |
| L Middle Temporal Gyrus | 1391.66 | 0.019 | -64 | -50 | 8 |
| L Lingual Gyrus | 1625.92 | 0.049 | -12 | -76 | -12 |
| R Superior Temporal Gyrus | 1533.82 | 0.020 | 56 | -44 | 12 |
| R Superior Frontal Gyrus | 1380.98 | 0.030 | 18 | 58 | 2 |
| R ParaHippocampal Gyrus | 1354.32 | 0.014 | 16 | -4 | -22 |
| L Middle Occipital Gyrus | 1211.79 | 0.044 | -24 | -74 | 24 |
| R Superior Medial Gyrus | 1164.52 | 0.044 | 12 | 54 | 0 |
| R Postcentral Gyrus | 1107.50 | 0.030 | 62 | -18 | 34 |
| Undefined | 941.03 | 0.044 | 42 | -44 | 18 |
| R Middle Frontal Gyrus | 903.57 | 0.050 | 26 | 46 | 6 |
| R Temporal Pole | 854.79 | 0.036 | 34 | 14 | -30 |
| L Superior Medial Gyrus | 854.40 | 0.045 | -16 | 58 | -2 |
| R Heschls Gyrus | 836.60 | 0.048 | 50 | -8 | 4 |
| L PCC | 821.29 | 0.047 | -8 | -44 | 20 |
| L PCC | 817.28 | 0.049 | -6 | -44 | 10 |
| R Area Fo3 | 807.00 | 0.028 | 22 | 32 | -12 |
| L Amygdala | 779.50 | 0.047 | -24 | -4 | -24 |
| Undefined | 685.16 | 0.041 | 4 | 0 | -18 |
| R Thal: Prefrontal | 648.55 | 0.044 | 6 | -6 | -4 |

**Table S4**. Blind subjects: Emotional facial expression sounds > object sounds. Only local maxima that are more than 24 mm apart are reported. TFCE - threshold free cluster enhancement; FDR - false discovery rate; MNI - Montreal Neurological Institute; Anatomical localization of MNI coordinates was performed using Anatomy SPM toolbox (v.2.2c).

| Region | TFCE value | p(FDR-corrected) | x | y | z (MNI space) |
|---|---|---|---|---|---|
| R Heschls Gyrus | 25980.93 | 0.001 | 54 | -12 | 6 |

| | | | | | |
|---|---|---|---|---|---|
| L Superior Temporal Gyrus | 25691.57 | 0.001 | -54 | -16 | 6 |
| L Middle Temporal Gyrus | 9334.78 | 0.001 | -62 | -42 | 10 |
| R Middle Temporal Gyrus | 7104.07 | 0.001 | 58 | -40 | 8 |
| R Temporal Pole | 4073.23 | 0.001 | 30 | 16 | -32 |
| L Temporal Pole | 3684.46 | 0.001 | -58 | 6 | -8 |
| R Superior Temporal Gyrus | 3240.63 | 0.001 | 44 | -2 | -14 |
| R IFG p. Orbitalis | 2775.78 | 0.001 | 48 | 24 | -12 |
| Undefined | 2416.92 | 0.001 | -34 | 2 | -20 |
| R Hippocampus | 1713.01 | 0.001 | 20 | -6 | -12 |
| L Temporal Pole | 1212.8 | 0.001 | -38 | 26 | -22 |
| R IFG p. Triangularis | 1048.87 | 0.001 | 48 | 22 | 24 |
| L Precentral Gyrus | 960.23 | 0.001 | -50 | -4 | 50 |
| L Thalamus | 715.29 | 0.001 | -2 | -16 | -2 |
| L Insula Lobe | 871.08 | 0.002 | -34 | -22 | 22 |
| Undefined | 708.7 | 0.002 | 18 | 30 | -14 |
| L Mid Orbital Gyrus | 536.63 | 0.002 | -6 | 48 | -14 |
| L IFG p. Triangularis | 697.39 | 0.003 | -48 | 22 | 12 |
| Undefined | 547.59 | 0.003 | 34 | 4 | 8 |
| L Olfactory cortex | 543.71 | 0.003 | -4 | 26 | -4 |
| L Postcentral Gyrus | 542.19 | 0.003 | -60 | -2 | 28 |
| R Inferior Occipital Gyrus | 886.2 | 0.004 | 42 | -78 | -8 |
| R Middle Temporal Gyrus | 708.7 | 0.004 | 44 | -60 | 10 |
| L Hippocampus | 620.79 | 0.004 | -26 | -24 | -10 |
| R Precentral Gyrus | 682.52 | 0.006 | 52 | -4 | 46 |
| R Fusiform Gyrus | 655.1 | 0.008 | 40 | -48 | -18 |
| R Lingual Gyrus | 687.74 | 0.015 | 16 | -62 | 0 |
| L Calcarine Gyrus | 623.97 | 0.015 | 4 | -88 | 8 |
| L Cerebelum Crus 2 | 417.38 | 0.015 | -10 | -82 | -42 |
| L Middle Occipital Gyrus | 587.56 | 0.016 | -42 | -82 | -2 |
| R Superior Frontal Gyrus | 237.57 | 0.016 | 36 | -4 | 66 |
| L Lingual Gyrus | 615.07 | 0.017 | -18 | -62 | -4 |
| L Cerebelum VIII | 339.31 | 0.018 | -26 | -64 | -48 |
| R Precuneus | 383.85 | 0.019 | 10 | -52 | 26 |
| R ParaHippocampal Gyrus | 276.42 | 0.027 | 16 | -30 | -14 |
| Undefined | 159.46 | 0.029 | 8 | -24 | -16 |
| R Superior Medial Gyrus | 301.5 | 0.031 | 8 | 56 | 28 |
| R Lingual Gyrus | 419.72 | 0.033 | 20 | -90 | -10 |
| R Postcentral Gyrus | 221.57 | 0.04 | 44 | -26 | 60 |
| R Superior Occipital Gyrus | 276.42 | 0.042 | 26 | -84 | 20 |
| L Cerebelum VI | 225.86 | 0.043 | -22 | -62 | -22 |
| L Superior Occipital Gyrus | 220.28 | 0.043 | -10 | -102 | 16 |
| R Precentral Gyrus | 275.66 | 0.048 | 36 | -16 | 40 |
| R Posterior-Medial Frontal | 170.29 | 0.05 | 6 | 6 | 66 |

**Table S5**. Blind subjects: Non-emotional facial expression sounds > object sounds. Only local maxima that are more than 24 mm apart are reported. TFCE - threshold free cluster enhancement; FDR - false discovery rate; MNI - Montreal Neurological Institute; Anatomical localization of MNI coordinates was performed using Anatomy SPM toolbox (v.2.2c).

| Region | TFCE value | p(FDR-corrected) | x | y | z (MNI) |
|---|---|---|---|---|---|
| R Superior Temporal Gyrus | 6383.23 | 0.001 | 54 | -20 | -4 |
| R Middle Temporal Gyrus | 4849.49 | 0.001 | 56 | -42 | 10 |
| L Middle Temporal Gyrus | 4385.69 | 0.001 | -60 | -20 | -2 |
| L Middle Temporal Gyrus | 3975.82 | 0.001 | -58 | -42 | 10 |
| R Temporal Pole | 2033.68 | 0.001 | 56 | 10 | -12 |
| R IFG p. Triangularis | 1527.14 | 0.001 | 52 | 26 | 8 |
| L IFG p. Triangularis | 1317.15 | 0.001 | -40 | 28 | -2 |
| R Posterior-Medial Frontal | 1807.07 | 0.002 | 8 | 2 | 66 |
| L Precentral Gyrus | 1480.12 | 0.002 | -48 | -4 | 50 |
| R Precentral Gyrus | 1384.88 | 0.002 | 50 | -2 | 44 |
| R Inferior Occipital Gyrus | 1351.39 | 0.002 | 48 | -76 | -4 |
| R Insula Lobe | 1134.4 | 0.002 | 36 | 4 | 8 |
| R Fusiform Gyrus | 1522.75 | 0.003 | 38 | -60 | -20 |
| L Middle Occipital Gyrus | 1350.77 | 0.003 | -50 | -74 | 2 |
| L Temporal Pole | 1073.03 | 0.003 | -54 | 10 | -14 |
| R Inferior Temporal Gyrus | 811.84 | 0.003 | 38 | 0 | -44 |
| R SupraMarginal Gyrus | 1212.02 | 0.004 | 58 | -18 | 26 |
| L Fusiform Gyrus | 1145.39 | 0.004 | -42 | -42 | -22 |
| L Precentral Gyrus | 1118.51 | 0.004 | -60 | 2 | 28 |
| L Insula Lobe | 986.75 | 0.004 | -34 | 4 | 10 |
| R Lingual Gyrus | 1056.41 | 0.005 | 16 | -64 | 0 |
| R IFG p. Orbitalis | 698.56 | 0.005 | 34 | 20 | -24 |
| R Cuneus | 1272.67 | 0.006 | 6 | -86 | 14 |
| R Superior Frontal Gyrus | 916.54 | 0.006 | 34 | 0 | 64 |
| L Cerebelum VI | 1138.28 | 0.008 | -26 | -62 | -24 |
| L Calcarine Gyrus | 994.96 | 0.008 | -12 | -68 | 10 |
| R Lingual Gyrus | 905.14 | 0.009 | 20 | -94 | -8 |
| L Calcarine Gyrus | 721.4 | 0.009 | -12 | -96 | -2 |
| L Superior Frontal Gyrus | 872.05 | 0.01 | -24 | -4 | 52 |
| R Middle Frontal Gyrus | 726 | 0.01 | 28 | 48 | 18 |
| L Middle Frontal Gyrus | 700.02 | 0.01 | -32 | 52 | 22 |
| R MCC | 937.05 | 0.011 | 8 | 18 | 36 |
| L Inferior Occipital Gyrus | 552.38 | 0.011 | -34 | -90 | -10 |
| L Putamen | 645.6 | 0.012 | -18 | 16 | -10 |
| L ACC | 695.43 | 0.013 | -6 | 34 | 24 |
| R MCC | 597.44 | 0.013 | 10 | -22 | 44 |
| L Cerebelum VII | 513.89 | 0.013 | -8 | -76 | -44 |
| Area PFm IPL | 270.92 | 0.023 | -56 | -52 | 46 |
| L Cuneus | 660.27 | 0.025 | -6 | -84 | 38 |

| | | | | | |
|---|---|---|---|---|---|
| L Cerebelum VIII | 554.49 | 0.029 | -30 | -64 | -48 |
| R Superior Medial Gyrus | 302.7 | 0.029 | 8 | 64 | 24 |
| R Thalamus | 232.72 | 0.03 | 14 | -20 | 8 |
| Undefined | 357.21 | 0.034 | -16 | -28 | 60 |
| L Middle Frontal Gyrus | 428.2 | 0.034 | -30 | 28 | 34 |
| Area Id1 | 233.8 | 0.036 | -36 | -18 | -6 |
| R Precuneus | 488.19 | 0.037 | 14 | -72 | 46 |
| R Cerebelum Crus 2 | 216.83 | 0.041 | 12 | -78 | -40 |
| R Mid Orbital Gyrus | 206.03 | 0.044 | 6 | 48 | -14 |
| R Cerebelum IX | 201.16 | 0.046 | 10 | -58 | -46 |
| R Rectal Gyrus | 224.21 | 0.046 | 4 | 30 | -16 |
| L Precuneus | 343.23 | 0.047 | -12 | -56 | 56 |
| L Paracentral Lobule | 371.63 | 0.047 | -8 | -24 | 60 |
| L Thalamus | 221.64 | 0.049 | -10 | -22 | 2 |
| R SupraMarginal Gyrus | 319.38 | 0.049 | 62 | -46 | 38 |

**Table S6**. Blind subjects: Speech sounds > object sounds. Only local maxima that are more than 24 mm apart are reported. TFCE - threshold free cluster enhancement; FDR - false discovery rate; MNI - Montreal Neurological Institute; Anatomical localization of MNI coordinates was performed using Anatomy SPM toolbox (v.2.2c).

| Region | TFCE value | p(FDR-corrected) | x | y | z (MNI) |
|---|---|---|---|---|---|
| R Precuneus | 1092.67 | 0.001 | 4 | -56 | 20 |
| R Heschls Gyrus | 10307.04 | 0.001 | 54 | -10 | 6 |
| L Superior Temporal Gyrus | 19057.35 | 0.001 | -54 | -16 | 6 |
| L Temporal Pole | 4369.28 | 0.001 | -56 | 6 | -6 |
| R Superior Temporal Gyrus | 1490.65 | 0.001 | 48 | -38 | 6 |
| L Precentral Gyrus | 1035.38 | 0.002 | -50 | -4 | 46 |
| R Inferior Temporal Gyrus | 572.88 | 0.002 | 38 | 0 | -42 |
| L Temporal Pole | 702.69 | 0.003 | -42 | 22 | -22 |
| R Middle Temporal Gyrus | 356.15 | 0.004 | 68 | -40 | -8 |
| Undefined | 856.56 | 0.008 | 44 | 16 | -16 |
| R Mid Orbital Gyrus | 564.04 | 0.009 | 4 | 62 | -4 |
| L Postcentral Gyrus | 467.7 | 0.009 | -34 | -22 | 42 |
| L SupraMarginal Gyrus | 811.72 | 0.015 | -46 | -44 | 24 |
| R Inferior Temporal Gyrus | 358.46 | 0.019 | 54 | -14 | -24 |
| R Precentral Gyrus | 822.33 | 0.02 | 50 | -8 | 44 |
| R Angular Gyrus | 225.7 | 0.022 | 54 | -62 | 26 |
| L Precentral Gyrus | 467.7 | 0.024 | -62 | 8 | 20 |
| R Rectal Gyrus | 528.59 | 0.024 | 4 | 38 | -18 |
| R Superior Medial Gyrus | 259.3 | 0.028 | 14 | 50 | 16 |
| R Precuneus | 305.32 | 0.037 | 16 | -40 | 4 |
| R Superior Frontal Gyrus | 249.98 | 0.046 | 16 | 32 | 44 |
| R Superior Frontal Gyrus | 255.19 | 0.047 | 16 | 60 | 26 |

| | | | | | |
|---|---|---|---|---|---|
| R Superior Frontal Gyrus | 248.17 | 0.048 | 20 | 50 | 40 |
| L Superior Medial Gyrus | 252.52 | 0.05 | -6 | 58 | 24 |

**Table S7**. Blind subjects: Animal sounds > object sounds. Only local maxima that are more than 24 mm apart are reported. TFCE - threshold free cluster enhancement; FDR - false discovery rate; MNI - Montreal Neurological Institute; Anatomical localization of MNI coordinates was performed using Anatomy SPM toolbox (v.2.2c).

| Region | TFCE value | p(FDR-corrected) | x | y | z (MNI) |
|---|---|---|---|---|---|
| L Superior Temporal Gyrus | 12157.63 | 0.001 | -54 | -16 | 6 |
| R Heschls Gyrus | 12445.18 | 0.001 | 52 | -12 | 6 |
| R Insula Lobe | 2303.67 | 0.001 | 42 | 2 | -12 |
| R IFG p. Orbitalis | 1079.83 | 0.001 | 32 | 30 | -12 |
| L Temporal Pole | 3140.26 | 0.001 | -56 | 6 | -6 |
| Undefined | 1484.36 | 0.001 | -36 | 2 | -20 |
| L Insula Lobe | 1256.25 | 0.001 | -32 | -14 | 18 |
| L IFG p. Orbitalis | 906.64 | 0.001 | -34 | 30 | -16 |
| R Middle Temporal Gyrus | 1329.35 | 0.002 | 60 | -38 | 6 |
| R Medial Temporal Pole | 652.63 | 0.004 | 32 | 6 | -34 |
| Undefined | 733.67 | 0.004 | -48 | -38 | -2 |
| L ACC | 1025.64 | 0.005 | -6 | 42 | 12 |
| R Superior Frontal Gyrus | 745.53 | 0.005 | 16 | 62 | 16 |
| L Rectal Gyrus | 921.32 | 0.006 | -6 | 36 | -18 |
| R ACC | 543.66 | 0.006 | 4 | 22 | 22 |
| R Middle Temporal Gyrus | 531.01 | 0.006 | 62 | -16 | -20 |
| R Mid Orbital Gyrus | 910.93 | 0.007 | 2 | 60 | -4 |
| L Superior Frontal Gyrus | 841.6 | 0.009 | -20 | 44 | 44 |
| L Superior Frontal Gyrus | 818.85 | 0.009 | -20 | 64 | 10 |
| L Middle Temporal Gyrus | 813.19 | 0.01 | -66 | -30 | -16 |
| L Postcentral Gyrus | 478.79 | 0.01 | -34 | -18 | 44 |
| L Superior Medial Gyrus | 751.73 | 0.011 | -8 | 26 | 62 |
| L Superior Temporal Gyrus | 297.24 | 0.016 | -58 | -44 | 22 |
| Undefined | 381.13 | 0.017 | -30 | -20 | -10 |
| L IFG p. Triangularis | 335.54 | 0.019 | -48 | 26 | 8 |
| R Olfactory cortex | 162.76 | 0.02 | 8 | 18 | -10 |
| L Middle Frontal Gyrus | 255.45 | 0.021 | -36 | 50 | 26 |
| R IFG p. Triangularis | 218.29 | 0.022 | 44 | 32 | 2 |
| R MCC | 446.51 | 0.023 | 10 | -12 | 50 |
| R Cerebelum IV-V | 514.41 | 0.024 | 12 | -62 | -10 |
| L Postcentral Gyrus | 498.23 | 0.025 | -60 | -12 | 42 |
| R Calcarine Gyrus | 252.87 | 0.025 | 30 | -62 | 8 |
| R Precuneus | 520.03 | 0.026 | 10 | -54 | 28 |
| R Calcarine Gyrus | 275.75 | 0.027 | 22 | -94 | -4 |
| R Superior Orbital Gyrus | 386.02 | 0.028 | 26 | 54 | -6 |

| | | | | | |
|---|---|---|---|---|---|
| L Lingual Gyrus | 446.41 | 0.03 | -16 | -60 | -6 |
| R Precuneus | 442.34 | 0.032 | 12 | -40 | 6 |
| R Middle Frontal Gyrus | 251.01 | 0.034 | 32 | 38 | 20 |
| L Paracentral Lobule | 474.82 | 0.038 | -8 | -26 | 62 |
| R Precentral Gyrus | 321.39 | 0.039 | 42 | -12 | 64 |
| R Precentral Gyrus | 303.01 | 0.041 | 52 | -4 | 44 |
| R Postcentral Gyrus | 417.25 | 0.042 | 16 | -30 | 62 |
| L MCC | 481.69 | 0.043 | 0 | -12 | 38 |
| Undefined | 164.51 | 0.044 | -30 | 14 | 28 |
| L Calcarine Gyrus | 378.26 | 0.045 | -2 | -76 | 14 |
| L Angular Gyrus | 224.36 | 0.047 | -50 | -72 | 24 |
| R MCC | 363.47 | 0.048 | 14 | -22 | 40 |
| L Fusiform Gyrus | 251.65 | 0.048 | -42 | -42 | -20 |
| L Calcarine Gyrus | 370.34 | 0.048 | -8 | -86 | 10 |
| R Posterior-Medial Frontal | 231.68 | 0.048 | 10 | 20 | 64 |
| R Superior Medial Gyrus | 163.52 | 0.048 | 4 | 54 | 42 |
| L Cuneus | 297.64 | 0.048 | -18 | -56 | 22 |
| L MCC | 405.93 | 0.049 | -6 | -4 | 36 |
| R Inferior Temporal Gyrus | 268.79 | 0.049 | 44 | -74 | -8 |
| R Superior Frontal Gyrus | 301.41 | 0.049 | 16 | 32 | 42 |
| L Caudate Nucleus | 180.06 | 0.05 | -18 | 24 | 2 |
| R Fusiform Gyrus | 271.14 | 0.05 | 40 | -48 | -22 |

**Table S8**. Blind subjects: Emotional expression sounds > speech sounds. Only local maxima that are more than 24 mm apart are reported. TFCE - threshold free cluster enhancement; FDR - false discovery rate; MNI - Montreal Neurological Institute; Anatomical localization of MNI coordinates was performed using Anatomy SPM toolbox (v.2.2c).

| Region | TFCE value | p(FDR-corrected) | x | y | z (MNI) |
|---|---|---|---|---|---|
| R Superior Temporal Gyrus | 3690.1 | 0.001 | 60 | -24 | 8 |
| R IFG p. Triangularis | 2612.74 | 0.001 | 52 | 28 | 24 |
| R IFG p. Opercularis | 2236.77 | 0.001 | 42 | 8 | 34 |
| Area Id1 | 2184.03 | 0.001 | 40 | -8 | -12 |
| R Inferior Occipital Gyrus | 2044.68 | 0.001 | 42 | -76 | -8 |
| R Fusiform Gyrus | 1821.85 | 0.001 | 38 | -48 | -14 |
| R Middle Temporal Gyrus | 1611.59 | 0.001 | 42 | -62 | 12 |
| L Superior Temporal Gyrus | 1564.87 | 0.001 | -42 | -18 | -2 |
| R Temporal Pole | 1543.45 | 0.001 | 42 | 18 | -26 |
| L Calcarine Gyrus | 1503.93 | 0.001 | 2 | -90 | 8 |
| R IFG p. Orbitalis | 1492.81 | 0.001 | 52 | 28 | -4 |
| L Cerebelum Crus 2 | 1492.17 | 0.001 | -8 | -80 | -42 |
| L Middle Occipital Gyrus | 1480.42 | 0.001 | -44 | -74 | 6 |
| L Middle Temporal Gyrus | 1379.7 | 0.001 | -64 | -50 | 8 |
| R Lingual Gyrus | 1282.77 | 0.001 | 14 | -62 | 2 |

| | | | | | |
|---|---|---|---|---|---|
| Undefined | 1144.17 | 0.001 | -34 | 2 | -20 |
| L Cerebelum VIII | 1000.2 | 0.001 | -28 | -66 | -46 |
| Thal: Temporal | 899.23 | 0.001 | 4 | -8 | 2 |
| L IFG p. Triangularis | 876.81 | 0.001 | -56 | 20 | 16 |
| R Superior Occipital Gyrus | 852.53 | 0.001 | 24 | -80 | 16 |
| Undefined | 642.68 | 0.001 | -8 | -20 | -16 |
| L Lingual Gyrus | 1258.39 | 0.002 | -18 | -62 | -2 |
| L IFG p. Triangularis | 822.88 | 0.002 | -36 | 14 | 28 |
| L IFG p. Triangularis | 806.97 | 0.002 | -40 | 24 | -2 |
| L Fusiform Gyrus | 852.53 | 0.003 | -20 | -82 | -18 |
| Area 33 | 565.09 | 0.003 | 0 | 22 | -4 |
| L Cuneus | 710.99 | 0.004 | -10 | -96 | 28 |
| L Middle Orbital Gyrus | 507.93 | 0.004 | -46 | 48 | -6 |
| Undefined | 687.6 | 0.005 | 26 | 30 | -10 |
| L Middle Occipital Gyrus | 695.56 | 0.006 | -18 | -102 | 0 |
| Cerebellar Vermis 3 | 461.19 | 0.006 | -2 | -48 | -22 |
| L Fusiform Gyrus | 497.85 | 0.008 | -36 | -48 | -20 |
| R Cerebelum Crus 2 | 470.63 | 0.008 | 16 | -76 | -40 |
| L Insula Lobe | 324.67 | 0.008 | -34 | -22 | 22 |
| Thal: Temporal | 448.34 | 0.016 | 22 | -32 | 6 |
| R Cuneus | 377.5 | 0.028 | 12 | -90 | 36 |
| R Middle Temporal Gyrus | 450.4 | 0.032 | 62 | -38 | -12 |
| Undefined | 148.88 | 0.036 | -22 | -36 | -34 |
| R Rolandic Operculum | 192.14 | 0.039 | 56 | 0 | 16 |
| R Cerebelum III | 231.94 | 0.044 | 16 | -32 | -22 |
| L Thalamus | 233.42 | 0.048 | -18 | -26 | 10 |

**Table S9**. Blind subjects: Non-emotional expression sounds > speech sounds. Only local maxima that are more than 24 mm apart are reported. TFCE - threshold free cluster enhancement; FDR - false discovery rate; MNI - Montreal Neurological Institute; Anatomical localization of MNI coordinates was performed using Anatomy SPM toolbox (v.2.2c).

| Region | TFCE value | p(FDR-corrected) | x | y | z (MNI) |
|---|---|---|---|---|---|
| R Precentral Gyrus | 3564.79 | 0.001 | 46 | 6 | 34 |
| R IFG p. Triangularis | 3243.98 | 0.001 | 50 | 32 | 10 |
| L Calcarine Gyrus | 2810.45 | 0.001 | -2 | -90 | 6 |
| Undefined | 2732.05 | 0.001 | 48 | -52 | 0 |
| R SupraMarginal Gyrus | 2500.87 | 0.001 | 64 | -38 | 24 |
| R Fusiform Gyrus | 2220.42 | 0.001 | 36 | -60 | -20 |
| L Middle Occipital Gyrus | 1909.46 | 0.001 | -48 | -72 | 2 |
| L Cerebelum VII | 1900.28 | 0.001 | -12 | -76 | -44 |
| R Superior Temporal Gyrus | 1845.67 | 0.001 | 50 | -22 | -4 |
| R Calcarine Gyrus | 1832.8 | 0.001 | 14 | -66 | 6 |
| L Cuneus | 1738.73 | 0.001 | -4 | -82 | 40 |

| | | | | | |
|---|---|---|---|---|---|
| R Fusiform Gyrus | 1712.2 | 0.001 | 24 | -84 | -12 |
| L Middle Temporal Gyrus | 1654.93 | 0.001 | -62 | -50 | 8 |
| L Cerebelum VI | 1627.11 | 0.001 | -38 | -66 | -20 |
| L Insula Lobe | 1489.91 | 0.001 | -32 | 16 | 4 |
| L Middle Occipital Gyrus | 1447.14 | 0.001 | -24 | -98 | 12 |
| L Cerebelum VI | 1367.95 | 0.001 | -10 | -78 | -18 |
| L Lingual Gyrus | 1363.67 | 0.001 | -20 | -62 | 2 |
| L IFG p. Opercularis | 1294.31 | 0.001 | -56 | 16 | 16 |
| L Precentral Gyrus | 1250.3 | 0.001 | -42 | 4 | 32 |
| R Middle Occipital Gyrus | 1196.58 | 0.001 | 26 | -86 | 14 |
| R Cerebelum Crus 2 | 1135.64 | 0.001 | 14 | -76 | -40 |
| R Inferior Parietal Lobule | 1023.45 | 0.001 | 32 | -54 | 48 |
| R Middle Frontal Gyrus | 1012.74 | 0.001 | 48 | 32 | 38 |
| L IFG p. Triangularis | 928.17 | 0.001 | -50 | 44 | 2 |
| L Pallidum | 577.49 | 0.001 | -22 | -6 | 4 |
| R Pallidum | 848.94 | 0.002 | 20 | 8 | 4 |
| L Middle Frontal Gyrus | 828.62 | 0.002 | -46 | 38 | 28 |
| R Superior Temporal Gyrus | 754.45 | 0.002 | 56 | 2 | -14 |
| L Medial Temporal Pole | 157.74 | 0.003 | -38 | 12 | -30 |
| Undefined | 219.41 | 0.006 | -8 | -40 | 22 |
| L Superior Medial Gyrus | 381.1 | 0.008 | 0 | 34 | 40 |
| Cerebellar Vermis 4/5 | 379.47 | 0.008 | -2 | -52 | -22 |
| Undefined | 261.77 | 0.009 | 24 | 28 | -10 |
| R Middle Frontal Gyrus | 533.05 | 0.01 | 40 | 58 | 8 |
| L Inferior Parietal Lobule | 334.86 | 0.011 | -52 | -56 | 48 |
| R Posterior-Medial Frontal | 657.73 | 0.012 | 8 | 12 | 56 |
| R MCC | 489.6 | 0.012 | 2 | 2 | 30 |
| R Thalamus | 342.83 | 0.012 | 10 | -22 | 6 |
| L Inferior Temporal Gyrus | 482.81 | 0.014 | -48 | -44 | -22 |
| R ACC | 406.18 | 0.014 | 6 | 24 | 16 |
| L Posterior-Medial Frontal | 570.86 | 0.015 | -10 | -6 | 66 |
| L Middle Temporal Gyrus | 128.51 | 0.016 | -54 | -22 | -8 |
| Undefined | 660.46 | 0.017 | -28 | -46 | 40 |
| Undefined | 399.68 | 0.017 | 22 | -8 | 48 |
| L MCC | 447.96 | 0.018 | -8 | -22 | 46 |
| L Cerebelum IV-V | 391.06 | 0.018 | -26 | -38 | -32 |
| L Superior Parietal Lobule | 543.4 | 0.02 | -24 | -68 | 50 |
| R Fusiform Gyrus | 212.04 | 0.021 | 42 | -18 | -26 |
| R Cerebelum IX | 192.6 | 0.021 | 14 | -50 | -40 |
| L SupraMarginal Gyrus | 449.64 | 0.023 | -56 | -28 | 30 |
| R SupraMarginal Gyrus | 424.97 | 0.023 | 48 | -34 | 42 |
| Undefined | 238.72 | 0.024 | 26 | 36 | 24 |
| R Superior Medial Gyrus | 193.4 | 0.025 | 6 | 60 | 30 |
| L Olfactory cortex | 169.85 | 0.033 | -18 | 6 | -18 |
| L Area Id1 | 157.9 | 0.045 | -38 | -20 | -6 |

| | | | | | |
|---|---|---|---|---|---|
| R ACC | 129.36 | 0.049 | 8 | 42 | 18 |
| R Precuneus | 59.82 | 0.049 | 20 | -62 | 30 |
| Undefined | 114.09 | 0.05 | -4 | -24 | -18 |

**Table S10**. Blind subjects: Emotional expression sounds > animal sounds. Only local maxima that are more than 24 mm apart are reported. TFCE - threshold free cluster enhancement; FDR - false discovery rate; MNI - Montreal Neurological Institute; Anatomical localization of MNI coordinates was performed using Anatomy SPM toolbox (v.2.2c).

| Region | TFCE value | p(FDR-corrected) | x | y | z (MNI) |
|---|---|---|---|---|---|
| L Middle Temporal Gyrus | 3538.34 | 0.002 | -60 | -20 | 0 |
| R Superior Temporal Gyrus | 4369.64 | 0.002 | 56 | -20 | -2 |
| R Middle Temporal Gyrus | 2064.63 | 0.002 | 56 | -42 | 10 |
| R Temporal Pole | 1505.78 | 0.002 | 52 | 10 | -14 |
| R Amygdala | 1173.13 | 0.002 | 22 | -6 | -14 |
| L Middle Temporal Gyrus | 1725.31 | 0.002 | -64 | -44 | 8 |
| R Middle Frontal Gyrus | 730.65 | 0.004 | 50 | -4 | 56 |
| R Temporal Pole | 932.85 | 0.005 | 30 | 16 | -32 |
| R IFG p. Triangularis | 854.43 | 0.005 | 44 | 24 | 22 |
| R Caudate Nucleus | 324.1 | 0.005 | 16 | 10 | 6 |
| R Olfactory cortex | 280.27 | 0.007 | 4 | 18 | -14 |
| R Precentral Gyrus | 571.41 | 0.008 | 56 | 2 | 18 |
| R Inferior Occipital Gyrus | 908.92 | 0.01 | 40 | -78 | -8 |
| L Hippocampus | 513.99 | 0.012 | -20 | -8 | -12 |
| L Superior Temporal Gyrus | 180.56 | 0.015 | -40 | -34 | 12 |
| R IFG p. Triangularis | 632.22 | 0.02 | 54 | 38 | 2 |
| Undefined | 283.81 | 0.021 | -4 | -2 | -14 |
| R Fusiform Gyrus | 639.61 | 0.021 | 40 | -46 | -14 |
| Undefined | 171.94 | 0.032 | 0 | -20 | -12 |
| L Precentral Gyrus | 243.96 | 0.033 | -50 | -2 | 52 |
| Undefined | 232.96 | 0.046 | 6 | -28 | -8 |
| L Middle Occipital Gyrus | 406.58 | 0.048 | -44 | -82 | -2 |
| L Inferior Occipital Gyrus | 348.54 | 0.049 | -38 | -78 | -8 |

**Table S11**. Blind subjects: Non-emotional expression sounds > animal sounds. Only local maxima that are more than 24 mm apart are reported. TFCE - threshold free cluster enhancement; FDR - false discovery rate; MNI - Montreal Neurological Institute; Anatomical localization of MNI coordinates was performed using Anatomy SPM toolbox (v.2.2c).

| Region | TFCE value | p(FDR-corrected) | x | y | z (MNI) |
|---|---|---|---|---|---|
| R Superior Temporal Gyrus | 1592.87 | 0.013 | 56 | -44 | 12 |
| R Superior Temporal Gyrus | 1386.91 | 0.013 | 52 | -20 | -6 |
| Undefined | 653.98 | 0.016 | -30 | 14 | 4 |
| L IFG p. Opercularis | 625.85 | 0.016 | -54 | 14 | 16 |

| | | | | | |
|---|---|---|---|---|---|
| L Middle Temporal Gyrus | 951.16 | 0.017 | -52 | -48 | 10 |
| L Posterior-Medial Frontal | 860.41 | 0.017 | -4 | 0 | 66 |
| L Precentral Gyrus | 605.05 | 0.017 | -44 | -2 | 52 |
| R IFG p. Triangularis | 665.6 | 0.019 | 54 | 36 | 6 |
| L Middle Temporal Gyrus | 489.83 | 0.019 | -58 | -20 | -4 |
| R IFG p. Opercularis | 970.27 | 0.02 | 48 | 18 | 26 |
| R Middle Temporal Gyrus | 947.35 | 0.02 | 48 | -66 | 6 |
| R Temporal Pole | 375.36 | 0.02 | 54 | 8 | -16 |
| R Middle Frontal Gyrus | 822.09 | 0.02 | 42 | 6 | 54 |
| R Insula Lobe | 626.63 | 0.024 | 32 | 22 | 4 |
| R Inferior Temporal Gyrus | 506.65 | 0.025 | 46 | -42 | -18 |
| L Inferior Temporal Gyrus | 326.96 | 0.029 | -46 | -40 | -22 |
| L Middle Frontal Gyrus | 324.52 | 0.035 | -28 | 44 | 18 |
| Undefined | 255.78 | 0.037 | -12 | 8 | 34 |
| R Middle Frontal Gyrus | 344.03 | 0.037 | 30 | 48 | 18 |
| L IFG p. Triangularis | 239.71 | 0.047 | -38 | 38 | 8 |
| R Posterior-Medial Frontal | 407.05 | 0.048 | 4 | 22 | 50 |
| L Inferior Parietal Lobule | 490.55 | 0.049 | -44 | -44 | 38 |
| R Cerebelum VI | 545.06 | 0.05 | 28 | -60 | -22 |
| L Cerebelum VI | 480.36 | 0.05 | -26 | -56 | -30 |
| R SupraMarginal Gyrus | 399.35 | 0.05 | 66 | -38 | 36 |

**Table S12**. Sighted subjects: Emotional facial expression sounds > object sounds. Only local maxima that are more than 24 mm apart are reported. TFCE - threshold free cluster enhancement; FDR - false discovery rate; MNI - Montreal Neurological Institute; Anatomical localization of MNI coordinates was performed using Anatomy SPM toolbox (v.2.2c).

| Region | TFCE value | p(FDR-corrected) | x | y | z (MNI) |
|---|---|---|---|---|---|
| L Superior Temporal Gyrus | 34208.38 | 0.001 | -54 | -16 | 6 |
| R Heschls Gyrus | 52631.74 | 0.001 | 54 | -10 | 6 |
| R Temporal Pole | 6024.31 | 0.001 | 54 | 14 | -12 |
| L Temporal Pole | 2062.56 | 0.001 | -54 | 14 | -12 |
| R Temporal Pole | 1066.78 | 0.006 | 38 | 24 | -28 |
| L Middle Occipital Gyrus | 618.01 | 0.006 | -36 | -92 | 10 |
| L Superior Occipital Gyrus | 666.83 | 0.011 | -12 | -102 | 12 |
| R Superior Occipital Gyrus | 487.66 | 0.011 | 20 | -96 | 20 |
| L Temporal Pole | 699.92 | 0.011 | -30 | 18 | -32 |
| R Middle Temporal Gyrus | 472.07 | 0.024 | 66 | -52 | -2 |
| R Precentral Gyrus | 613.5 | 0.025 | 54 | -6 | 46 |
| L Precentral Gyrus | 297.15 | 0.036 | -50 | -4 | 54 |
| R ACC | 288.53 | 0.042 | 10 | 38 | -2 |
| L Postcentral Gyrus | 202.11 | 0.044 | -56 | -10 | 40 |
| R Middle Occipital Gyrus | 306.4 | 0.045 | 40 | -88 | 0 |
| L Inferior Occipital Gyrus | 256.59 | 0.047 | -36 | -70 | -8 |

| | | | | | |
|---|---|---|---|---|---|
| L Superior Occipital Gyrus | 331.07 | 0.047 | -20 | -90 | 34 |
| R Lingual Gyrus | 178.46 | 0.048 | 16 | -76 | -10 |

**Table S13**. Sighted subjects: Non-emotional facial expression sounds > object sounds. Only local maxima that are more than 24 mm apart are reported. TFCE - threshold free cluster enhancement; FDR - false discovery rate; MNI - Montreal Neurological Institute; Anatomical localization of MNI coordinates was performed using Anatomy SPM toolbox (v.2.2c).

| Region | TFCE value | p(FDR-corrected) | x | y | z (MNI) |
|---|---|---|---|---|---|
| L Superior Temporal Gyrus | 3230.28 | 0.002 | -60 | -16 | 2 |
| R Superior Temporal Gyrus | 7557.82 | 0.002 | 64 | -22 | 4 |
| R Temporal Pole | 2718.02 | 0.002 | 54 | 10 | -14 |
| R Postcentral Gyrus | 637.63 | 0.002 | 62 | -2 | 20 |
| L Middle Temporal Gyrus | 1790.07 | 0.002 | -62 | -40 | 6 |
| L Postcentral Gyrus | 1545.16 | 0.002 | -62 | 0 | 20 |
| L Temporal Pole | 906.06 | 0.006 | -56 | 6 | -16 |
| R Middle Temporal Gyrus | 906 | 0.025 | 66 | -52 | -2 |
| R Inferior Temporal Gyrus | 691.39 | 0.028 | 38 | 4 | -42 |
| R ACC | 339.18 | 0.031 | 2 | 28 | -4 |
| L ParaHippocampal Gyrus | 426.22 | 0.032 | -26 | -4 | -28 |
| L Medial Temporal Pole | 355.08 | 0.039 | -32 | 20 | -36 |
| L IFG p. Triangularis | 637.92 | 0.045 | -46 | 26 | 2 |
| R Rectal Gyrus | 334.39 | 0.048 | 12 | 16 | -16 |
| L Superior Temporal Gyrus | 390.88 | 0.05 | -50 | -38 | 20 |

**Table S14**. Sighted subjects: Speech sounds > object sounds. Only local maxima that are more than 24 mm apart are reported. TFCE - threshold free cluster enhancement; FDR - false discovery rate; MNI - Montreal Neurological Institute; Anatomical localization of MNI coordinates was performed using Anatomy SPM toolbox (v.2.2c).

| Region | TFCE value | p(FDR-corrected) | x | y | z (MNI) |
|---|---|---|---|---|---|
| R Heschls Gyrus | 27466.74 | 0.002 | 54 | -10 | 6 |
| L Superior Temporal Gyrus | 22248.32 | 0.002 | -58 | -14 | 4 |
| L Superior Temporal Gyrus | 3271.94 | 0.002 | -48 | -36 | 18 |
| L Temporal Pole | 1790.13 | 0.002 | -54 | 12 | -10 |
| L Postcentral Gyrus | 1378.63 | 0.002 | -48 | -8 | 54 |
| R Middle Temporal Gyrus | 1721.74 | 0.003 | 66 | -8 | -18 |
| R Temporal Pole | 1383.53 | 0.003 | 52 | 16 | -16 |
| R ACC | 578.54 | 0.019 | 10 | 40 | -2 |
| R Temporal Pole | 478.69 | 0.031 | 34 | 22 | -32 |
| R Cuneus | 780.05 | 0.031 | 18 | -100 | 12 |
| R Middle Temporal Gyrus | 335.09 | 0.035 | 68 | -40 | 0 |
| R Thalamus | 347.52 | 0.037 | 16 | -26 | 0 |
| R Superior Frontal Gyrus | 265.08 | 0.037 | 14 | 58 | 36 |

| | | | | | |
|---|---|---|---|---|---|
| L PCC | 719.95 | 0.037 | 2 | -52 | 28 |
| L Superior Occipital Gyrus | 1117.33 | 0.039 | -16 | -94 | 26 |
| R Superior Occipital Gyrus | 1090.52 | 0.039 | 20 | -82 | 32 |
| R Lingual Gyrus | 996.33 | 0.04 | 20 | -66 | -12 |
| R Middle Occipital Gyrus | 959.86 | 0.04 | 40 | -86 | 2 |
| L Cuneus | 872.65 | 0.04 | -16 | -70 | 20 |
| L Precuneus | 195.15 | 0.041 | -4 | -64 | 48 |
| R Fusiform Gyrus | 594.05 | 0.042 | 36 | -42 | -16 |
| R Calcarine Gyrus | 952.69 | 0.042 | 24 | -64 | 14 |
| L Middle Occipital Gyrus | 937.09 | 0.043 | -36 | -90 | 10 |
| L Medial Temporal Pole | 299.51 | 0.044 | -48 | 16 | -28 |
| R Precentral Gyrus | 512.96 | 0.044 | 58 | -6 | 44 |
| L Superior Medial Gyrus | 323.05 | 0.044 | -14 | 50 | -2 |
| R Superior Frontal Gyrus | 220.71 | 0.046 | 18 | 40 | 52 |
| L Superior Medial Gyrus | 335.63 | 0.047 | -4 | 56 | 12 |
| R Angular Gyrus | 416.2 | 0.047 | 50 | -60 | 24 |
| L Lingual Gyrus | 669.62 | 0.048 | -22 | -46 | -6 |
| R Superior Medial Gyrus | 331.59 | 0.05 | 2 | 56 | 14 |
| L Lingual Gyrus | 788.64 | 0.05 | -18 | -56 | -8 |

**Table S15**. Sighted subjects: Animal sounds > object sounds. Only local maxima that are more than 24 mm apart are reported. TFCE - threshold free cluster enhancement; FDR - false discovery rate; MNI - Montreal Neurological Institute; Anatomical localization of MNI coordinates was performed using Anatomy SPM toolbox (v.2.2c).

| Region | TFCE value | p(FDR-corrected) | x | y | z (MNI) |
|---|---|---|---|---|---|
| R Heschls Gyrus | 16800.8 | 0.002 | 54 | -10 | 6 |
| L Heschls Gyrus | 13259.77 | 0.002 | -50 | -18 | 8 |
| R Insula Lobe | 594.31 | 0.017 | 40 | 6 | -14 |
| L Amygdala | 341.24 | 0.021 | -20 | -4 | -20 |
| L Temporal Pole | 519.93 | 0.039 | -46 | 8 | -12 |
| R Cuneus | 376.38 | 0.049 | 10 | -92 | 28 |

**Table S16**. Sighted subjects: Emotional expression sounds > speech sounds. Only local maxima that are more than 24 mm apart are reported. TFCE - threshold free cluster enhancement; FDR - false discovery rate; MNI - Montreal Neurological Institute; Anatomical localization of MNI coordinates was performed using Anatomy SPM toolbox (v.2.2c).

| Region | TFCE value | p(FDR-corrected) | x | y | z (MNI) |
|---|---|---|---|---|---|
| R Superior Temporal Gyrus | 2924.31 | 0.003 | 54 | -26 | 10 |
| R Area Id1 | 2092.02 | 0.003 | 40 | -8 | -12 |
| L Superior Temporal Gyrus | 1750.18 | 0.003 | -42 | -20 | -4 |
| L Cerebelum Crus 2 | 1182.05 | 0.003 | -16 | -78 | -36 |
| L Superior Temporal Gyrus | 1069.02 | 0.009 | -38 | -38 | 16 |

| | | | | | |
|---|---|---|---|---|---|
| R IFG p. Triangularis | 1129.52 | 0.011 | 42 | 18 | 24 |
| R Middle Temporal Gyrus | 315.82 | 0.029 | 66 | -50 | 2 |
| R IFG p. Triangularis | 666.09 | 0.034 | 46 | 28 | 0 |
| Undefined | 857.35 | 0.036 | -32 | 2 | -18 |
| Undefined | 464.32 | 0.036 | -10 | -2 | -12 |
| R IFG p. Orbitalis | 445.75 | 0.04 | 38 | 20 | -22 |
| R IFG p. Orbitalis | 469.64 | 0.041 | 40 | 20 | -14 |
| R Insula Lobe | 550.4 | 0.043 | 28 | 24 | -8 |
| L IFG p. Triangularis | 405.28 | 0.044 | -34 | 36 | 8 |
| Cerebellar Vermis 7 | 379.47 | 0.045 | 0 | -78 | -24 |
| L Temporal Pole | 270.24 | 0.046 | -34 | 8 | -28 |
| L IFG p. Triangularis | 465.73 | 0.047 | -36 | 26 | 10 |
| R Middle Frontal Gyrus | 527.71 | 0.049 | 46 | 4 | 56 |
| R Precentral Gyrus | 524.99 | 0.05 | 48 | 4 | 46 |
| R Insula Lobe | 469.64 | 0.05 | 26 | 20 | -16 |

**Table S17**. Sighted subjects: Non-emotional expression sounds > speech sounds. Only local maxima that are more than 24 mm apart are reported. TFCE - threshold free cluster enhancement; FDR - false discovery rate; MNI - Montreal Neurological Institute; Anatomical localization of MNI coordinates was performed using Anatomy SPM toolbox (v.2.2c).

| Region | TFCE value | p(FDR-corrected) | x | y | z (MNI) |
|---|---|---|---|---|---|
| R Middle Temporal Gyrus | 1135 | 0.007 | 60 | -40 | 8 |
| L Cerebelum Crus 2 | 1423.05 | 0.007 | -16 | -78 | -36 |
| R Middle Frontal Gyrus | 1330.03 | 0.007 | 38 | 8 | 36 |
| R IFG p. Triangularis | 1079.5 | 0.007 | 50 | 26 | 24 |
| R IFG p. Triangularis | 1164.71 | 0.008 | 48 | 26 | -2 |
| R Middle Frontal Gyrus | 865.41 | 0.008 | 40 | 2 | 60 |
| R Middle Temporal Gyrus | 786.37 | 0.008 | 48 | -20 | -8 |
| R Cerebelum Crus 1 | 753.46 | 0.009 | 18 | -78 | -34 |
| L IFG p. Triangularis | 843.82 | 0.011 | -40 | 24 | 2 |
| L ParaHippocampal Gyrus | 530.61 | 0.013 | -24 | -4 | -28 |
| L Cerebelum VII | 528.69 | 0.019 | -34 | -64 | -48 |
| R ParaHippocampal Gyrus | 302.72 | 0.02 | 30 | 6 | -26 |
| L Putamen | 558.13 | 0.023 | -22 | 6 | -6 |
| R Posterior-Medial Frontal | 732.43 | 0.024 | 2 | 18 | 62 |
| Undefined | 192.97 | 0.026 | 8 | -16 | -14 |
| R Putamen | 298.16 | 0.03 | 26 | -2 | 6 |
| R Cerebelum Crus 1 | 387.99 | 0.03 | 42 | -76 | -32 |
| L Precentral Gyrus | 715.35 | 0.032 | -38 | 2 | 42 |
| R Medial Temporal Pole | 444.8 | 0.033 | 54 | 4 | -16 |
| R Putamen | 425.85 | 0.039 | 26 | 10 | 8 |
| R Rectal Gyrus | 192.32 | 0.042 | 12 | 22 | -16 |
| R Cerebelum VIII | 295.79 | 0.043 | 8 | -64 | -34 |

| | | | | | |
|---|---|---|---|---|---|
| L Superior Medial Gyrus | 391.69 | 0.046 | 0 | 44 | 40 |
| L IFG p. Opercularis | 616.46 | 0.046 | -42 | 16 | 20 |
| R Inferior Temporal Gyrus | 291.49 | 0.047 | 38 | 2 | -44 |
| Undefined | 180.08 | 0.048 | 4 | -28 | -12 |
| R Superior Frontal Gyrus | 168.92 | 0.048 | 16 | 56 | 22 |
| R Inferior Parietal Lobule | 230.73 | 0.048 | 56 | -52 | 42 |
| L Middle Temporal Gyrus | 269.78 | 0.049 | -62 | -44 | 6 |
| R Thalamus | 269.81 | 0.049 | 8 | -16 | 14 |
| Undefined | 213.87 | 0.049 | 32 | 40 | 16 |

**Table S18**. Sighted subjects: Emotional expression sounds > animal sounds. Only local maxima that are more than 24 mm apart are reported. TFCE - threshold free cluster enhancement; FDR - false discovery rate; MNI - Montreal Neurological Institute; Anatomical localization of MNI coordinates was performed using Anatomy SPM toolbox (v.2.2c).

| Region | TFCE value | p(FDR-corrected) | x | y | z (MNI) |
|---|---|---|---|---|---|
| L Cerebelum Crus 2 | 1224.75 | 0.002 | -18 | -78 | -36 |
| L Superior Temporal Gyrus | 6110.3 | 0.002 | -60 | -16 | 2 |
| R Superior Temporal Gyrus | 13586.46 | 0.002 | 58 | -12 | -2 |
| R Temporal Pole | 1183.6 | 0.002 | 38 | 24 | -28 |
| L Middle Temporal Gyrus | 1365.37 | 0.002 | -62 | -42 | 8 |
| L Temporal Pole | 1107.1 | 0.002 | -56 | 4 | -12 |
| R Middle Temporal Gyrus | 1240.94 | 0.003 | 66 | -52 | -2 |
| R Middle Frontal Gyrus | 830.12 | 0.003 | 50 | -2 | 54 |
| R Area Id1 | 768.74 | 0.011 | 46 | 0 | -20 |
| R Insula Lobe | 896.82 | 0.013 | 40 | -14 | 14 |
| R Middle Frontal Gyrus | 699.51 | 0.013 | 34 | 16 | 42 |
| R IFG p. Triangularis | 858.61 | 0.016 | 48 | 18 | 22 |
| R Superior Medial Gyrus | 489.92 | 0.017 | 12 | 46 | 36 |
| L Precentral Gyrus | 259.52 | 0.019 | -48 | -4 | 52 |
| R IFG p. Triangularis | 876.52 | 0.021 | 48 | 28 | 0 |
| L ParaHippocampal Gyrus | 576.51 | 0.022 | -26 | 0 | -30 |
| R Superior Medial Gyrus | 411.87 | 0.029 | 14 | 54 | 2 |
| R IFG p. Opercularis | 370.55 | 0.03 | 42 | 10 | 8 |
| R Middle Frontal Gyrus | 369.12 | 0.03 | 26 | 26 | 34 |
| L Hippocampus | 376.79 | 0.035 | -26 | -14 | -12 |
| L Postcentral Gyrus | 432.8 | 0.04 | -44 | -12 | 34 |
| L ACC | 185.29 | 0.041 | -8 | 26 | 30 |
| R Cerebelum Crus 1 | 279.88 | 0.043 | 18 | -76 | -32 |
| R ACC | 159.06 | 0.047 | 6 | 32 | -4 |
| L IFG p. Triangularis | 341.75 | 0.048 | -48 | 28 | 2 |
| L Insula Lobe | 343.72 | 0.048 | -30 | 18 | -16 |

**Table S19**. Sighted subjects: Non-emotional expression sounds > animal sounds. Only local maxima that are more than 24 mm apart are reported. TFCE - threshold free cluster enhancement; FDR - false discovery rate; MNI - Montreal Neurological Institute; Anatomical localization of MNI coordinates was performed using Anatomy SPM toolbox (v.2.2c).

| Region | TFCE value | p(FDR-corrected) | x | y | z (MNI) |
|---|---|---|---|---|---|
| L Cerebelum Crus 2 | 1212.08 | 0.006 | -18 | -78 | -34 |
| R Middle Temporal Gyrus | 2827.85 | 0.006 | 58 | -34 | 4 |
| R Temporal Pole | 869.78 | 0.014 | 54 | 8 | -16 |
| L Middle Temporal Gyrus | 576 | 0.014 | -62 | -42 | 6 |
| L Superior Medial Gyrus | 294.96 | 0.023 | -12 | 36 | 28 |
| R Middle Frontal Gyrus | 574.9 | 0.039 | 28 | 26 | 32 |
| R Temporal Pole | 415.52 | 0.044 | 40 | 16 | -28 |
| L Middle Temporal Gyrus | 675.77 | 0.045 | -56 | -22 | -6 |
| Undefined | 778.47 | 0.046 | 26 | 22 | -6 |
| R SupraMarginal Gyrus | 495.51 | 0.046 | 58 | -44 | 34 |
| R Superior Orbital Gyrus | 418.12 | 0.047 | 32 | 58 | -4 |
| R Inferior Temporal Gyrus | 526.31 | 0.049 | 38 | 2 | -44 |
| R IFG p. Triangularis | 974.8 | 0.049 | 48 | 28 | -2 |
| R Middle Frontal Gyrus | 1017.83 | 0.049 | 40 | 12 | 40 |
| R Superior Orbital Gyrus | 353.05 | 0.05 | 22 | 42 | -12 |
| R IFG p. Triangularis | 850.81 | 0.05 | 42 | 12 | 22 |

**Table S20**. Interaction analysis: (Blind subjects > sighted subjects) x (all expression sounds > all other sounds). Only local maxima that are more than 24 mm apart are reported. TFCE - threshold free cluster enhancement; FDR - false discovery rate; MNI - Montreal Neurological Institute; Anatomical localization of MNI coordinates was performed using Anatomy SPM toolbox (v.2.2c).

| Region | TFCE value | p(FDR-corrected) | x | y | z (MNI) |
|---|---|---|---|---|---|
| R Middle Temporal Gyrus | 1188.48 | 0.043 | 48 | -68 | 4 |
| R Inferior Temporal Gyrus | 1090.83 | 0.043 | 44 | -42 | -20 |
| L Calcarine Gyrus | 977.56 | 0.043 | 0 | -90 | 6 |
| R Lingual Gyrus | 920.05 | 0.043 | 14 | -64 | 4 |
| R Superior Temporal Gyrus | 852.47 | 0.043 | 56 | -44 | 12 |
| L Lingual Gyrus | 805.48 | 0.043 | -14 | -56 | -4 |
| R Fusiform Gyrus | 746.11 | 0.043 | 26 | -82 | -12 |
| L Middle Occipital Gyrus | 1093.06 | 0.043 | -48 | -74 | 4 |
| L Middle Temporal Gyrus | 626.64 | 0.043 | -52 | -46 | 12 |
